# HCF101 is a novel component of the CIA cytosolic iron-sulfur synthesis pathway in the human pathogen *Toxoplasma gondii*

**DOI:** 10.1101/2024.05.01.592079

**Authors:** Eléa A. Renaud, Ambre J.M. Maupin, Laurence Berry, Julie Bals, Yann Bordat, Vincent Demolombe, Valérie Rofidal, Florence Vignols, Sébastien Besteiro

## Abstract

Several key cellular functions depend on proteins harboring an iron-sulfur (Fe-S) cofactor. As these Fe-S proteins localize to several subcellular compartments, they require a dedicated machinery for cofactor assembly. For instance, in plants and algae there are Fe-S cluster synthesis pathways localizing to the cytosol, but also present in the mitochondrion and in the chloroplast, two organelles of endosymbiotic origin. *Toxoplasma gondii* is a plastid-bearing parasitic protist responsible for a pathology affecting humans and other warm-blooded vertebrates. We have characterized the *Toxoplasma* homologue of HCF101, originally identified in plants as a protein transferring Fe-S clusters to photosystem I subunits in the chloroplast. Contrarily to plants, we have shown that HCF101 does not localize to the plastid in parasites, but instead is an important component of the cytosolic Fe-S assembly (CIA) pathway which is vital for *Toxoplasma*. While the CIA pathway is widely conserved in eukaryotes, it is the first time the involvement of HCF101 in this pan-eukaryotic machinery is established. Moreover, as this protein is essential for parasite viability and absent from its mammalian hosts, it constitutes a novel and promising potential drug target.

## Introduction

Iron-sulfur (Fe-S) clusters are ancient inorganic cofactors of proteins which are essential in virtually all forms of life [1]. These cofactors are found in a variety of proteins involved in numerous electron transfer and metabolic reactions that support key housekeeping cellular functions like respiration, photosynthesis, DNA repair and replication, protein translation, RNA modifications, but also in regulatory proteins and environmental signals sensors [2,3]. The most common clusters, which in most Fe-S proteins function as electron transfer groups [4], are in the rhombic [2Fe-2S], cubane [4Fe-4S], or more rarely the intermediate [3Fe-4S] configuration. Assembly of these cofactors is a carefully controlled process in order to avoid cellular toxicity arising from the accumulation of free Fe and S, which are highly reactive. Different Fe-S cluster assembly pathways developed early in evolution to synthesize the clusters and can be found in archaea, bacteria, and then evolved to mitochondrial, plastidic and cytosolic iron-sulfur assembly machineries in eukaryotes [5,6]. Thus, eukaryotic Fe-S cluster assembly machineries found in organelles of endosymbiotic origin were inherited from Alphaproteobacteria (the ‘iron sulfur cluster’, or ISC, pathway in the mitochondrion) and Cyanobacteria (the ‘sulfur mobilization’, or SUF, pathway in plastids), while a specific machinery (‘cytosolic iron-sulfur cluster assembly’, or CIA) is used for the biogenesis of cytosolic and nuclear Fe-S proteins.

While the individual components of these Fe-S assembly machineries display some structural diversity, they involve similar biochemical steps [7]. Typically, a cysteine desulfurase will generate sulfur from L-cysteine, scaffold proteins and chaperones will provide a molecular platform allowing the assembly of Fe and S into a cluster, and finally carrier proteins will deliver the cluster to target apoproteins. This happens in a similar fashion but with a specific molecular machinery in the three subcellular compartments in which Fe-S cluster synthesis takes place in eukaryotes, with a main difference that the early step of CIA depends on the mitochondrial machinery. In fact, although de novo cluster assembly has been suggested to happen in the cytosol of mammalian cells [8], experimental evidence in a large variety of eukaryotes points to the requirement of mitochondrial ISC components, in particular the Nfs1 cysteine desulfurase, to generate a S precursor that will be translocated to the cytosol [9–11]. In mammals and yeast, P-loop NTPases NUBP1 and NUBP2 are the scaffolds for initial [4Fe-4S] Fe-S cluster assembly in the CIA system [12] and electrons are provided by DRE2, together with NDOR1 (ATR3 in plants) [13]. Noticeably, some eukaryotes like plants lack a NUBP2 homologue and the scaffolding complex is instead composed of a dimer of NBP35, the NUBP1 homologue [14]. NAR1, a protein containing two [4Fe-4S] clusters, then acts as the Fe-S carrier and associates with the CIA targeting complex (CTC) [15]. The CTC, that will recognize client apoproteins through direct interactions and mediate the insertion of the Fe-S cluster, typically comprises MET18/MMS19 (plant/human nomenclature), CIA1/CIAO1, and AE7/CIAO2B [16].

*Toxoplasma gondii* is a widespread obligate intracellular parasitic protist belonging to the phylum Apicomplexa. Infection is usually harmless to immunocompetent individuals, but can lead to severe life-threatening disease in developing fetuses and immunocompromised individuals [17]. This parasite contains two organelles of endosymbiotic origin: a mitochondrion and a relict plastid called apicoplast, which are both of high metabolic importance [18]. Consequently, like in plants and algae *T. gondii* harbors three distinct Fe-S cluster synthesis pathways: the CIA in the cytosol, ISC in the mitochondrion and SUF in the apicoplast [19–21]. The SUF pathway being essential for the survival of the parasite and absent from its mammalian host, its components and client proteins are particularly attractive as potential drug targets [22], which prompted us to look for Fe-S proteins absent from mammals but present in plant and apicomplexan parasites. Investigating the Fe-S cluster assembly components in protists, which are highly divergent phylogenetically from the canonical yeast or mammalian cell models, can bring interesting insights into the evolution of this molecular machinery [23,24].

Here, we report the characterization of the *T. gondii* homologue of the high-chlorophyll-fluorescence 101 (HCF101) protein, which is typically a plastid-associated Fe-S transfer protein in the plant *Arabidopsis thaliana* [25,26]. Our work shows that TgHCF101 is essential for parasite viability and we demonstrate for the first time its implication in the eukaryotic CIA pathway.

## Results

### TgHCF101 is a cytosolic protein

HCF101 is a putative P-loop containing nucleoside triphosphate hydrolase that belongs to the Mrp/NBP35 family (also called ApbC) of dimeric iron-sulfur carrier proteins that are found ubiquitously in all domains of life and thus likely appeared early during evolution [27,28]. Compared with NBP35, in addition to the nucleotide hydrolase domain HCF101 also contains a N-terminal DUF59/MIP18-family domain [29], as well as a C-terminal domain of unknown function (DUF971). We performed homology searches in the http://www.ToxoDB.org database to retrieve members of the Mrp/NPB35 family encoded by the *T. gondii* genome. Besides TgNBP35, that has an unusual mitochondrial localization in *T. gondii* [20,24], in line with previous studies [24,30] we identified TGGT1_318590 as a potential homologue of HCF101. A broader phylogenetic analysis that included members of the Mrp/NPB35 family from a wide variety of eukaryotes (S1 Table), revealed that this putative *T. gondii* homologue clearly segregates with members of the HCF101 group, clearly distinct from NBP35 homologues (Fig. 1A). Moreover, its primary sequence analysis confirmed a three domain-organization typical of HCF101 proteins, while NBP35 homologues mostly consist of the central ATPase domain (Fig. 1B). Of note, our analysis, in accordance with a previous study [24], confirmed the absence of a CFD1 homologue in Apicomplexa, contrarily to what has been suggested by a recent report [31]. In conclusion, TGGT1_318590 may be named TgHCF101.

**Figure 1.**
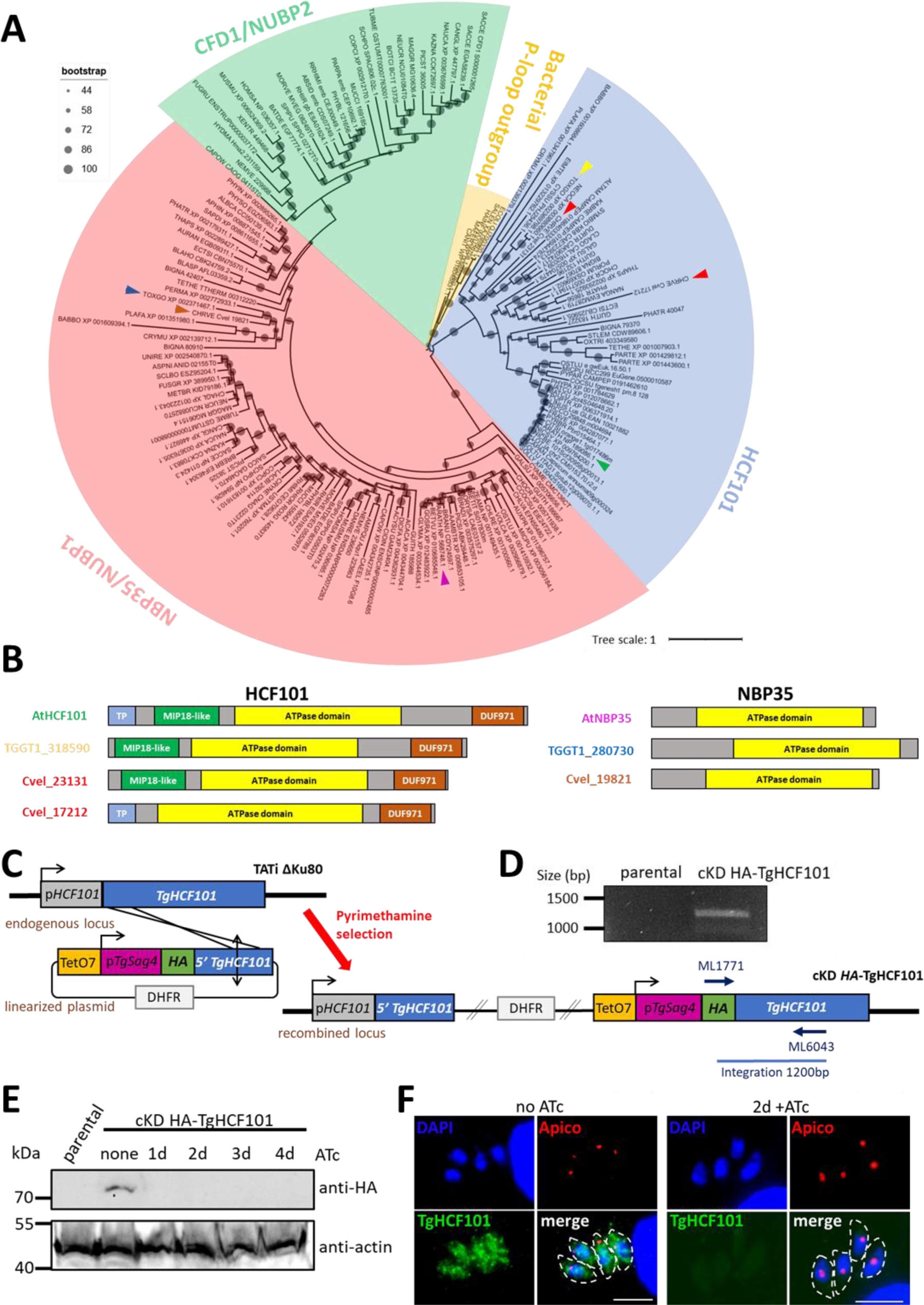
TgHCF101 is a cytosolic protein. **A.** Evolutionary relationship of proteins of the MRP family. Eukaryotic sequences from HCF101 homologs were aligned and submitted to phylogenetic analysis with the maximum likelihood method. Scale bar represents 1 residue substitution per site. Bacterial P-loop ATPase sequences were used as an outgroup. HCF101 from *A. thaliana* is indicated by a green arrowhead, TgHCF101 by a yellow arrowhead and the two homologs present in *C. velia* are indicated by red arrowheads. NBP35 homologs from *A. thaliana, T. gondii* and *C. velia* are indicated by purple, blue and brown arrowheads, respectively. **B.** Schematic representation of homologs for HCF101 in *A. thaliana* (AtHCF101), in *T. gondii* (TGGT1_318590) and in *C. velia* (Cvel_23131, Cvel_17212), and homologs for NBP35 in *A. thaliana* (AtNBP35), in *T. gondii* (TGGT1_280730) and in *C. velia* (Cvel_19821). Main domains are highlighted on the sequences; TP: predicted transit peptide, MIP18-like domain (or Domain of unknown function 59, DUF59), the ATPase domain and DUF971. **C.** Strategy for generating the inducible knockdown of *TgHCF101* by promoter replacement and simultaneous N-terminal tagging of the TgHCF101 protein in the TATi ΔKu80 cell line. **D.** Diagnostic PCR for checking correct integration using the primers mentioned in (B), on genomic DNAs of a transgenic parasite clone and of the parental strain. **E.** Immunoblot analysis of the cKD-TgHCF101 mutant and parental line showing efficient tagging and downregulation of TgHCF101 starting at 24h of treatment with ATc. Actin was used as a loading control. **F.** Immunofluorescence assay showing a cytosolic signal for TgHCF101 protein (labeled with an anti-HA antibody), with no particular co-localization with the apicoplast (Apico, labeled with anti-PDH-E2 antibody), and total depletion of the protein after 48h of ATc treatment. DNA was stained with 4′,6-diamidino-2-phenylindole (DAPI). Scale bar= 5 µm.

In *A. thaliana*, the *HCF101* mutant was found to be impaired in photosynthesis [32]: interfering with HCF101 function has an impact on the maturation of both photosystem I (PSI) and ferredoxin-thioredoxin reductases, which contain [4Fe-4S] clusters [25,26]. The apicoplast harbored by apicomplexan parasites has lost the ability to perform photosynthesis [33], and some members of the Apicomplexa phylum, like *Cryptosporidium*, have even completely lost the plastid [34] while it has retained a HCF101 homologue (Fig. 1A, S1 Fig.). Thus, a function linked to photosystem seems unlikely for apicomplexan parasites. Sequence analysis confirmed that, in contrast to plant homologues, TgHCF101 lacks a predicted N-terminal transit peptide, which is typically needed for plastid import (Fig. 1B). Interestingly, *Chromera velia*, a close non-parasitic relative of Apicomplexa whose plastid has retained its photosynthetic capacity [35], has two HCF101 isoforms: one with a predicted transit peptide (thus potentially addressed to the plastid), and one without (Fig. 1, S1 Fig.). Specific analysis of selected eukaryotic HCF101 homologs showed that Apicomplexa proteins, like ciliates and dinoflagellates, form distinct groups that are clearly separate from the Archaeplastida group of plants and algae bearing plastid-associated HCF101 isoforms (S1 Fig.). Interestingly, while the transit peptide-bearing *C. velia* isoform segregates with homologs of plastid-containing eukaryotes, the second isoform is closer to the Apicomplexa homologs (S1 Fig.). A recent study suggests that the plastid-located HCF101 likely evolved from an ancestral cytosolic version of the protein [24]. This led us to hypothesize that apicomplexan HCF101 would be involved in a cytosolic-related, rather than in a plastid-associated, Fe-S cluster transport function.

To assess experimentally the localization of TgHCF101, we first tagged an ectopic copy with a C-terminal GFP and observed a cytosolic localization (S2A Fig.), in line with what had been described previously when an additional copy of TgHCF101 bearing a C-terminal Human influenza hemagglutinin (HA) tag was expressed [24]. However, a stable TgHCF101-GFP cell line could not be established, suggesting the potential importance of C-terminal accessibility for proper protein function. We then next generated a transgenic cell line in which we modified the 5’ region of the *TgHCF101* gene by homologous recombination to replace the endogenous promoter by an inducible-Tet07*SAG4* promoter, and at the same time adding a sequence coding for a N-terminal hemagglutinin (HA) tag (Fig. 1C, D). In this conditional knock-down (cKD) HA-TgHCF101 cell line, the addition of anhydrotetracycline (ATc) can repress transcription of *TgHCF101* through a Tet-Off system [36]. Two independent transgenic clones were obtained and found to behave similarly in the initial phenotypic assays we performed, so only one will be described in details here.

Immunoblot analysis showed a N-terminal HA-tagged protein at around 70 kDa (the predicted molecular mass for TgHCF101), and depletion of the protein was efficient upon addition of ATc (Fig. 1E). Immunofluorescence assay (IFA) showed a punctate cytosolic signal for TgHCF101, clearly distinct from the apicoplast, and efficiently depleted by the addition of ATc (Fig. 1F). Disruption of the apicoplast-located SUF pathway has a strong impact on the lipoylation of the E2 subunit of the pyruvate dehydrogenase (PDH-E2) by Fe-S-containing lipoate synthase LipA [19,22]. However, depletion of TgHCF101 did not seem to have an impact on the lipoylation profile of PDH-E2 (S2B Fig.). Disruption of the SUF pathway also typically impacts the apicoplast-located isoprenoid synthesis, which involves two Fe-S oxidoreductases (IspG and IspH) and has been shown to have downstream consequences on the glycosylation of proteins involved in parasite motility [22]. We thus assessed the gliding motility of the parasites after TgHCF101 depletion and found that it was only affected after four days of ATc treatment, which might be reflecting long-term impact of TgHCF101 depletion on parasite fitness rather than a direct consequence of apicoplast-related Fe-S cluster synthesis (S3A, B Fig.).

Altogether, our results indicate that TgHCF101 is not associated with the apicoplast-located SUF pathway, but localizes to the cytosol instead.

### TgHCF101 is essential for parasite growth

We next assessed the impact of TgHCF101 depletion on the parasite’s lytic cycle: we evaluated the capacity of TgHCF101-depleted tachyzoites (a highly replicative and highly invasive form responsible for the acute form of the disease caused by *T. gondii*) to generate lysis plaques. For this, we infected a monolayer of host cells in absence or continuous presence of ATc for 7 days (Fig. 2A, B). Depletion of TgHCF101 completely prevented plaque formation. While absence of plaques highlights a fitness problem for the parasites, it does not necessarily implies their death: for instance, in response to stress, tachyzoites can convert into a slow-growing and cyst-enclosed persisting stage called bradyzoites (associated with the chronic form of the disease), that can reactivate when they encounter more favourable conditions [37]. We thus performed a similar experiment, in which the ATc was washed out at the end of the 7-day incubation, and then incubated the parasites for an extra 7 days in the absence or presence of ATc before evaluating plaque formation (Fig. 2C). We did not observe any plaques after ATc removal, suggesting that the parasites were not able to recover after 7 days of TgHCF101 depletion, likely because they were dead.

**Figure 2.**
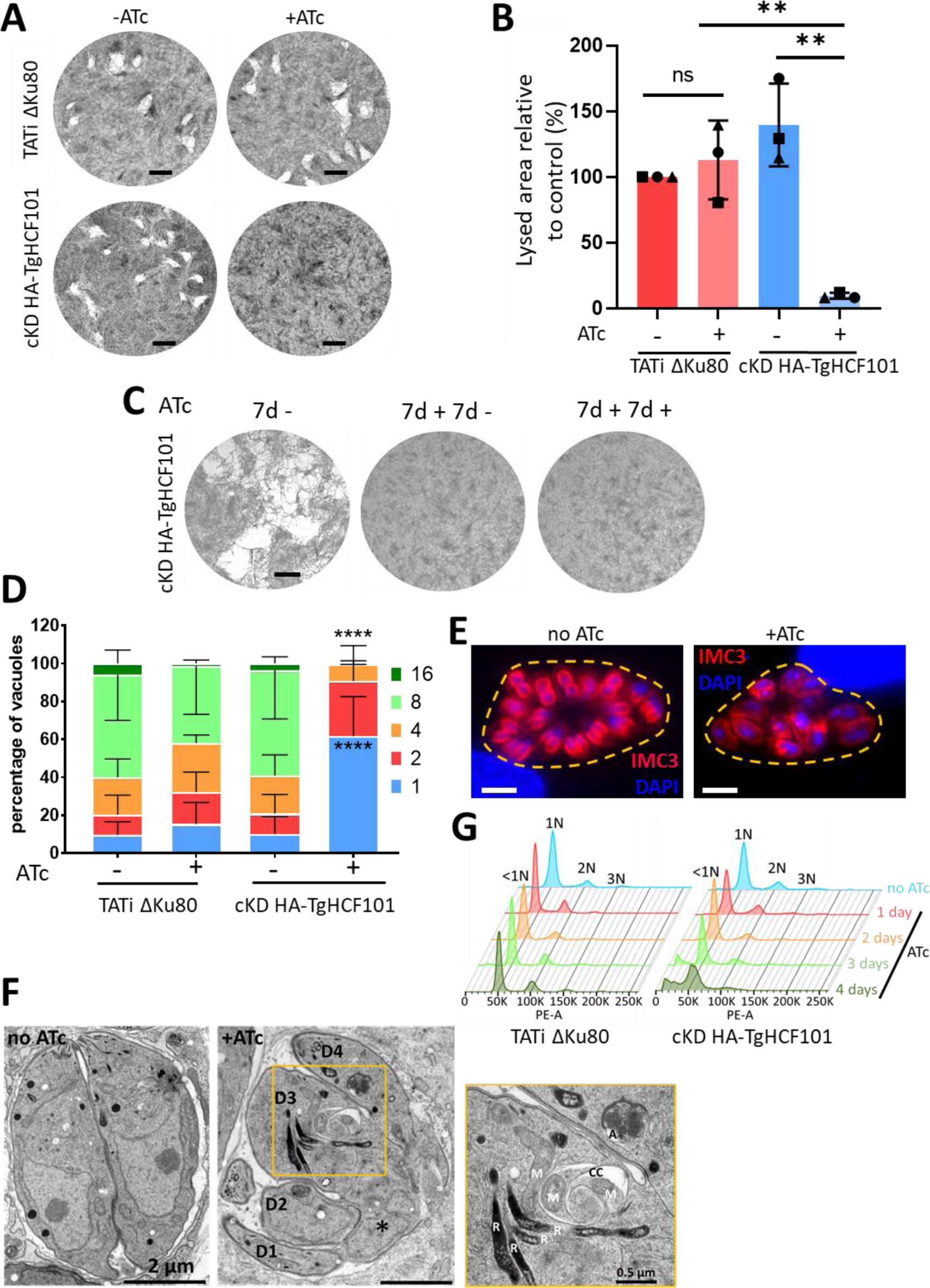
TgHCF101 is essential for parasite growth and survival. **A.** Plaque assays were carried out by infecting a monolayer of HFFs with TATi ΔKu80 or cKD-TgHCF101 cell lines for 7 days in the presence or absence of ATc. Scale bar= 2 mm. **B.** Quantification of plaque area observed in (A). Results are expressed as percentage of lysed area relative to control (TATi ΔKu80 -ATc, set as 100% for reference). Values represented are mean ± SD from n=3 independent biological replicates, ** p-value ≤ 0.01, Student *t*-test **C.** Plaque assays were carried out with cKD-TgHCF101 parasites as described in (A) but for the ‘7d+7d-‘ condition, ATc was washed-out after 7 days and parasites were allowed to grow for another 7 days without ATc, while in the ‘7d+7d+’ condition, ATc treatment was maintained for 7 more days. The ‘7d –‘ control was kept without ATc for 7 days of growth. Scale bar= 2mm. **D.** Replication assay of parental (TATi ΔKu80) and transgenic cell lines (cKD-TgHCF101): parasites were pre-cultured for 48h in the presence or absence of ATc and allowed to invade HFF-coated coverslips for another 24h in the presence or absence of ATc, for a total of up to 72h of treatment with ATc. Number of parasites per vacuole was quantified for each condition and expressed as a percentage, 200 vacuoles were counted for each condition. Values represented are mean ± SD of 3 independent biological replicates, **** p-value ≤ 0.0001 by two-way ANOVA with Dunnett’s multiple comparison test, showing a significant difference when comparing the TATi ΔKu80 control and cKD-TgHCF101+ATc parasites for percentage of vacuoles containing 1 or 8 parasites. **E.** DNA content analysis by flow cytometry on TATi ΔKu80 and cKD-TgHCF101 parasites treated or not with ATc up to 4 days and stained with propidium iodide. 1N, 2N, 3N represent the ploidy, with <1N corresponding to parasites with less than 1 full nuclear DNA content. **F.** Immunofluorescence assay of cKD-TgHCF101 parasites showing asynchronous division of parasites growing in the same vacuole (outlined with a yellow dotted line) upon TgHCF101 depletion (+ATc condition: parasites pre-incubated for 48h with ATc and allowed to invade coverslips for another 48h in the presence of ATc). Individual parasites and budding daughter cells are outlined by anti-IMC3 antibody staining (magenta), DNA was stained with DAPI. Scale bar= 5 µm. **G.** Electron microscopy of cKD-TgHCF101 parasites pre-incubated with ATc for 24h before being released from their host cell and allowed to reinvade for 24h in the presence (+ATc) or grown in absence of ATc (-ATc). D: daughter bud, R: rhoptry, A: apicoplast, M: mitochondrion, CC: cytoplasmic cleft. The asterisk denotes unincorporated nuclear material. Scale bar= 2 µm (0.5 µM for inset magnification).

To assess whether this defect in the lytic cycle is due to a replication problem, cKD HA-TgHCF101 parasites were preincubated in ATc for 48 hours and then released mechanically, before infecting new host cells for an additional 24 hours in ATc prior to parasite counting. We noted that incubation with ATc led to an accumulation of vacuoles with fewer parasites (Fig. 2D). To get more precise insights into the impact of TgHCF101 depletion on parasite division, we labeled dividing parasites with inner membrane complex (IMC) protein IMC3 [38] to detect growing daughter cells (Fig. 2E). *T. gondii* tachyzoites develop inside a mother cell by a process called endodyogeny [39] and this division is usually highly synchronous within the same parasitophorous vacuole, yet after two days of ATc treatment vacuoles showed a marked lack of synchronicity for daughter cell budding (Fig. 2E). Investigation of the consequences of TgHCF101 depletion on developing parasites by electron microscopy highlighted important defects on cytokinesis, including incomplete daughter cell budding or defaults in organellar segregation (Fig. 2F, S4 Fig.). We also observed occasional vacuoles and cytoplasmic clefts (Fig. 2F, S4 Fig.). However, the overall appearance of organelles seemed essentially normal (Fig. 2F, S4 Fig.). To get a more general overview, we performed IFA using organelle-specific markers, which confirmed that TgHCF101 depletion leads to defaults in the replication or segregation of organelles, some daughter cells appearing devoid of apicoplast, mitochondrial or nuclear material (S5 Fig.). Finally, through propidium iodide labeling and flow cytometer-based analysis of DNA content, we observed that long-term depletion of TgHCF101 led to the emergence of an increasing subpopulation of parasites with sub-1N DNA content, thus smaller than the typical haploid 1N DNA content of the parasites (Fig. 2G), suggesting an impairment in DNA synthesis.

Overall, our results indicate that TgHCF101 depletion leads to important and, apparently, irreversible defects in parasite replication and growth.

### Depletion of TgHCF101 induces stress response mechanisms

Through electron microscopy analysis, we also noticed that the depletion of TgHCF101 induced the appearance of structures resembling lipid droplets (LDs) in several parasites (Fig. 3A, S4 Fig.). In order to properly quantify this, we used Nile red, a selective fluorescent stain for neutral lipids, and microscopic imaging (Fig. 3B). We noticed a strong increase in both the number and size of LD in cKD HA-TgHCF101 parasites incubated for two days with ATc (Fig. 3C, D). To see if this was a common feature induced by abolishing Fe-S synthesis, we quantified LDs in mutants of the mitochondrial (TgISU1) and apicoplast (TgSUFC) pathways [19]. Depletion of these proteins did not lead to an increase in LD formation (S6A and B Fig.), indicating that LD induction may be linked to a pathway distinct from the mitochondrial or plastid-related Fe-S cluster assembly machinery. We then also tested a recently characterized mutant for TgABCB7L (a mitochondrial transporter of a sulfur precursor upstream of the CIA pathway [40]) and a mutant of the CIA shuttle protein TgNAR1 (www.ToxoDB.org entry TGGT1_242580) that we generated, and which is important for parasite fitness (S7 Fig.). Interestingly, both these CIA-related mutants induced an increase in LD numbers (although more modest for the TgABCB7L mutant), while only the TgNAR1 led to a statistically significant increase in LD size (Fig. 3C, D).

**Figure 3.**
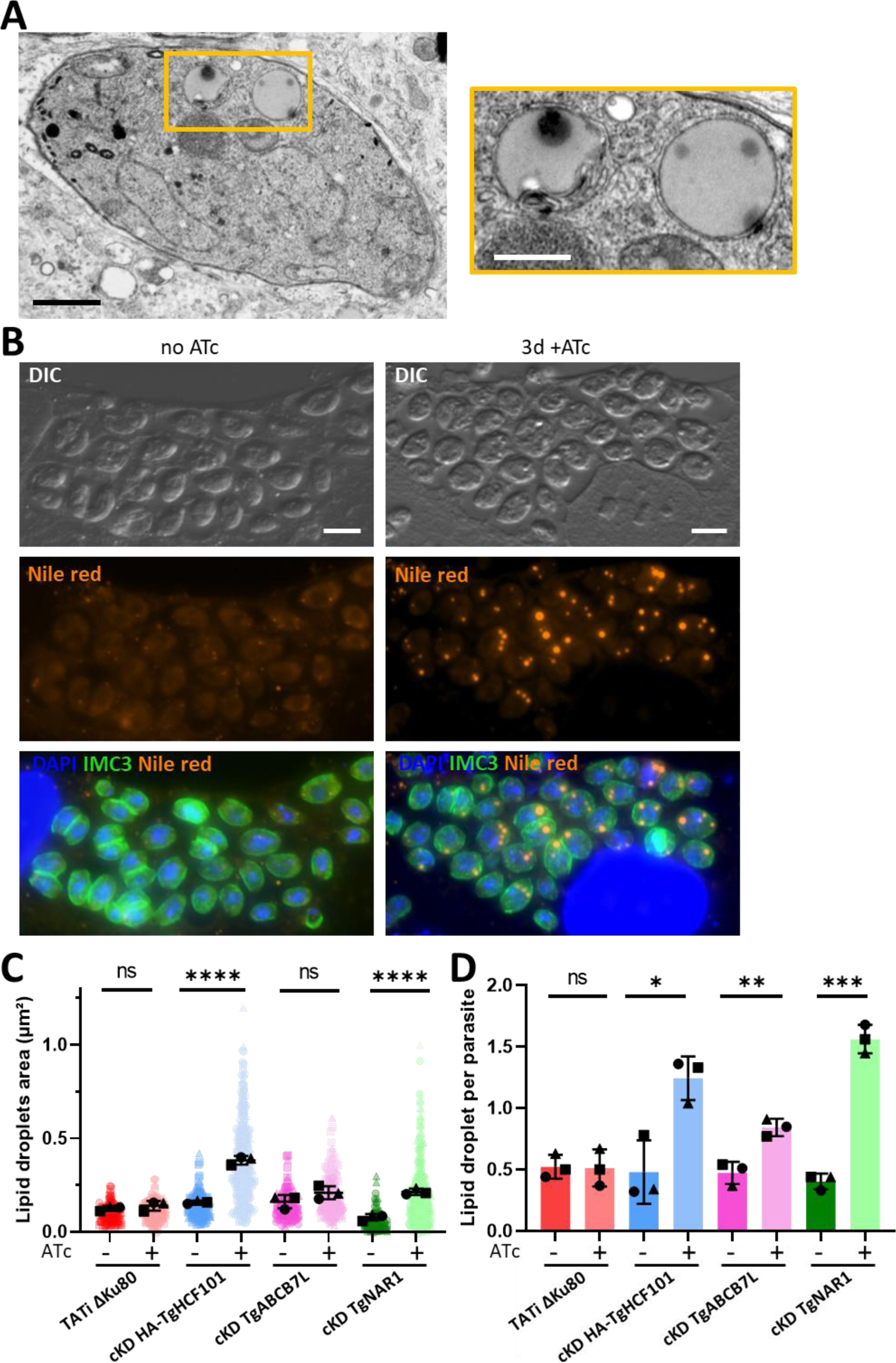
TgHCF101 depletion induces lipid droplet accumulation. **A.** Electron microscopy of cKD-TgHCF101 parasites pre-incubated with ATc for 48h and allowed to reinvade for 24h in the presence of ATc. Scale bar= 1 µm. The panel on the right corresponds to a magnification of the selection on the left panel, highlighting lipid droplets. Scale bar= 500 nm **B.** Fluorescent imaging of parasites from the cKD-TgHCF101 cell line treated for 72h with ATc or in grown the absence of ATc, shows an accumulation of lipid droplets upon TgHCF101 depletion. Lipid droplets were detected with Nile red (orange), parasites are outlined with an anti-IMC3 antibody (green) and DNA is stained with DAPI. DIC= Differential interference contrast. Scale bar= 5 µm. **C.** and **D.** correspond to the quantification of lipid droplet area and number, respectively. 100 parasites were analyzed per condition. The parental (TATi ΔKu80) and transgenic parasites (cKD HA-TgHCF101, cKD TgABCB7, cKD TgNAR1) were grown in absence of ATc, or in presence of ATc for 72h. Values are represented as the mean ± SD of n=3 independent biological replicates (different symbols represent different series); ns, not significant (p-value>0.05), * p-value ≤ 0.05, ** p-value ≤ 0.01, *** p-value ≤ 0.001, **** p-value ≤ 0.0001, Student’s *t*-test.

As LD droplet induction seems to be a feature common to CIA pathway mutants and to the TgHCF101 mutant, we thought that TgHCF101 may play a role in this pathway. Additional supporting evidence comes from the characterization of a C-terminal targeting complex recognition (TCR) signal ending with an aromatic residue ([LIM]-[DES]-[WF]) in the clients proteins of the cytosolic Fe-S cluster machinery or their adaptors [41], which can be clearly identified in the apicomplexan HCF101 homologues (Fig. 4A). Of note, this motif is also present in the *C. velia* isoform that has no transit peptide and thus which is likely to have a plastid-independent function (Fig. 4A). Importantly, the last three amino acids of TgHCF101 (LEW, Fig. 4A) constitute a canonical TCR motif. Thus, from the cKD HA-TgHCF101 mutant, we generated cell lines in which we expressed either a wild-type copy or one deleted for this putative TCR signal (Fig. 4B, C). As expected, expression of these additional copies was unaltered by ATc treatment (Fig. 4D), and the corresponding proteins localized to the cytoplasm (Fig. 4E). Upon depletion of the ATc-regulated copy, expressing the extra wild-type copy was efficient in restoring parasite growth (Fig. 4F, G) and preventing LD accumulation (Fig. 4H, I). Yet, in sharp contrast, expressing the TCR-deleted extra copy did not improve parasite fitness (Fig. 4F, G) and did not prevent induction of LD formation (Fig. 4H, I).

**Figure 4.**
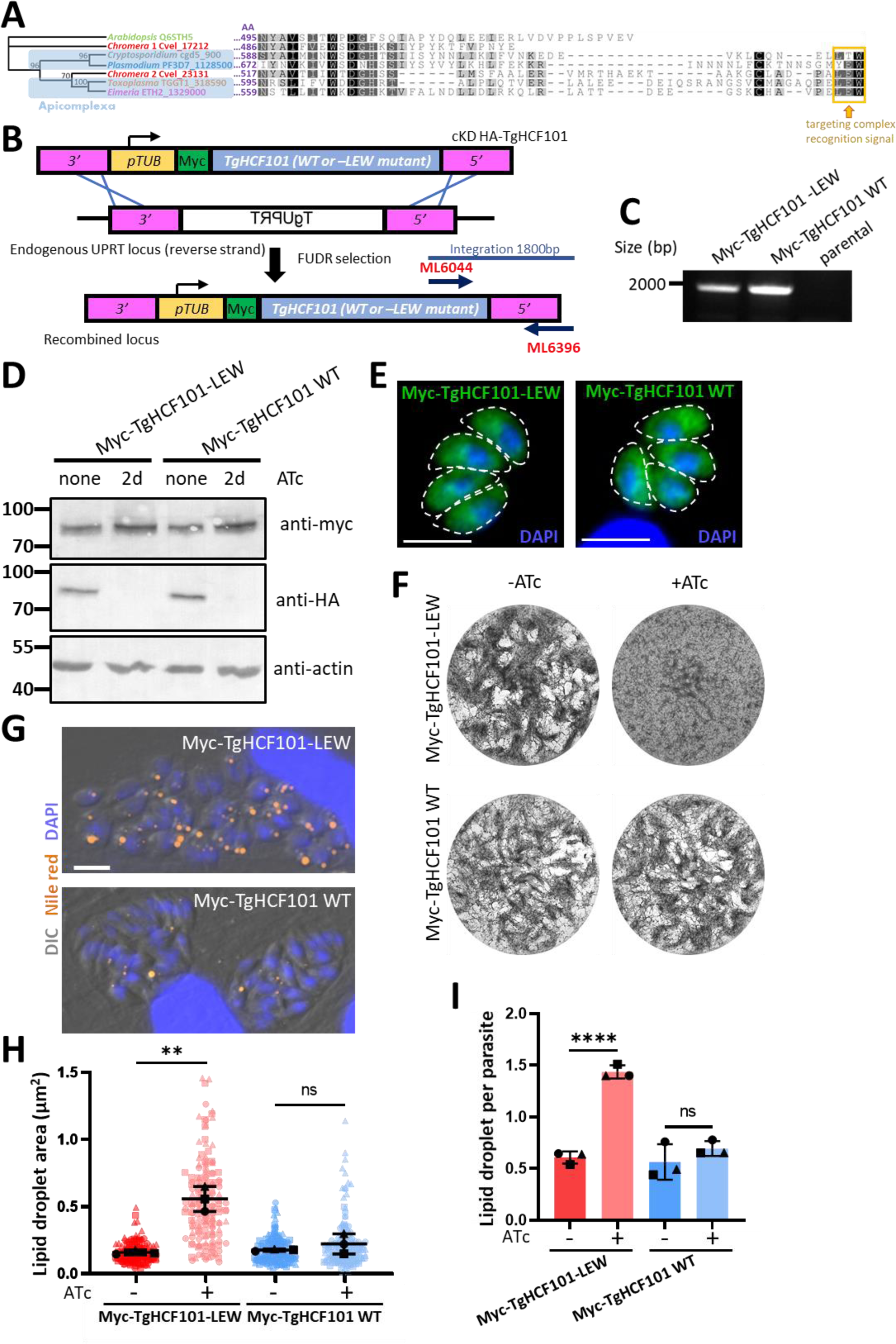
A C-terminal TCR-like motif is necessary for TgHCF101 function. **A**. Alignment of the C-terminal region of HCF101 homologs from different eukaryotes, including the plant *A. thaliana*, the two isoforms of Apicomplexa-relative photosynthetic algae *C. velia*, as well as several Apicomplexa species (*T. gondii*, *Eimeria tenella*, *P. falciparum* and apicoplast-less *Cryptosporidium parvum*). The tryptophan-containing targeting complex recognition (TCR) signal is highlighted in yellow. Consensus tree was obtained with protein alignment and bootstrap values (500 replicates) are indicated at the base of the nodes. **B.** Schematic representation of the strategy for generating cell lines expressing myc-tagged wild-type (WT) or TCR motif-deleted (-LEW) copies of TgHCF101 by integrating an extra copy of the gene of interest by double homologous recombination at the *Uracil Phosphoribosyltransferase* (*UPRT*) locus. Negative selection with 5-fluorodeoxyuridine (FUDR) was used to select transgenic parasites based on their absence of UPRT expression. **C.** Diagnostic PCR for verifying integration at the *UPRT* locus thanks to the primers described in **A. D.** Immunoblot analysis showing expression of the additional myc-tagged copies upon depletion of the ATc-regulated HA-tagged copy. **E.** Immunofluorescence with anti-myc antibody confirms the cytoplasmic localization of the extra copies. Parasite shape is outlined. DNA was stained with DAPI. Scale bar= 5 µm. **F.** Plaque assay showing restoration of growth by the additional WT copy upon depletion of the ATc-regulated copy, contrarily to the TCR mutant. **G.** Representative images of vacuoles (outlined) containing parasites grown continuously in the presence of ATc for 72 hours and stained by Nile red (orange) for lipid droplets. DIC: differential interference contrast. DNA was stained with DAPI. Scale bar= 10 µm. **H.** and **I.** correspond to the quantification of lipid droplet area and number, respectively. 100 parasites were analyzed per condition. WT or TCR-depleted (-LEW) parasites were grown in presence of ATc for 72h. Values are represented as the mean ± SD of n=3 independent biological replicates (different symbols represent different series); ns, not significant (p-value>0.05), ** p-value ≤ 0.01, **** p-value ≤ 0.0001, Student’s *t*-test.

To assess if the induction of LD formation could play a role in the demise of the parasites, we used T863, a pharmaceutical inhibitor of the acyl coenzyme A:diacylglycerol acyltransferase 1 (DGAT1) enzyme, which is important for storing neutral lipids in cytoplasmic LD [42]. While treatment with T863 efficiently decreased the number of LDs (S6C, D and E Fig.), long term incubation during plaque assays did not allow any recovery in parasite fitness (S6F Fig.). This suggests that LDs are not detrimental to TgHCF101-depleted parasites (or at least not solely responsible for their demise), but instead could be part of an integrated stress response, as they have been shown in other eukaryotes to be upregulated in response to cellular injuries including nutrient-related and oxidative stresses [43].

Disrupting proteins involved in Fe-S cluster assembly may have a direct effect on the stability and expression levels of local Fe-S proteins. Thus, to get insights into the effect of TgHCF101 depletion at the molecular level, we performed global label-free quantitative proteomic analyses. Of course, this may also affect downstream cellular pathways or functions, and other pathways may also be upregulated in compensation. cKD HA-TgHCF101 or TATi ΔKu80 parental control parasites were treated for three days with ATc prior to a global proteomic analysis and compared for protein expression. We selected candidates with a log_2_(fold change) ≤-0.55 or ≥0.55 (corresponding to a ∼1.47-fold change in decreased or increased expression) and a p-value <0.05 (ANOVA, *n*=4 biological replicates) and we completed this dataset by selecting some candidates that were consistently and specifically absent from the mutant cell lines or only expressed in these (S2 and S3 Tables). Many proteins with higher abundance were bradyzoite stage-specific and more particularly components of the cyst wall or their cell surface [44,45] (Fig. 5A, B, S2 Table). Tachyzoites can convert to the persistent bradyzoite form upon stress, so to verify if TgHCF101 depletion was inducing stage conversion we used a lectin from the plant *Dolichos biflorus*, which recognizes the SRS44/CST1 cyst wall glycoprotein in differentiating cysts [46] (Fig. 5C, D). We could see that TgHCF101 depletion induced the appearance of up to 10% of lectin-labeled vacuoles during the first 48 hours of intracellular growth in the presence of ATc. However, when parasites were kept for up to a week in the presence of ATc, the proportion of lectin-positive vacuoles plateaued at 14%, and only a very limited number of vacuoles looked like bona fide mature cysts (less than 2% were both lectin-positive and containing more than 2 parasites) (Fig. 5E).

**Figure 5.**
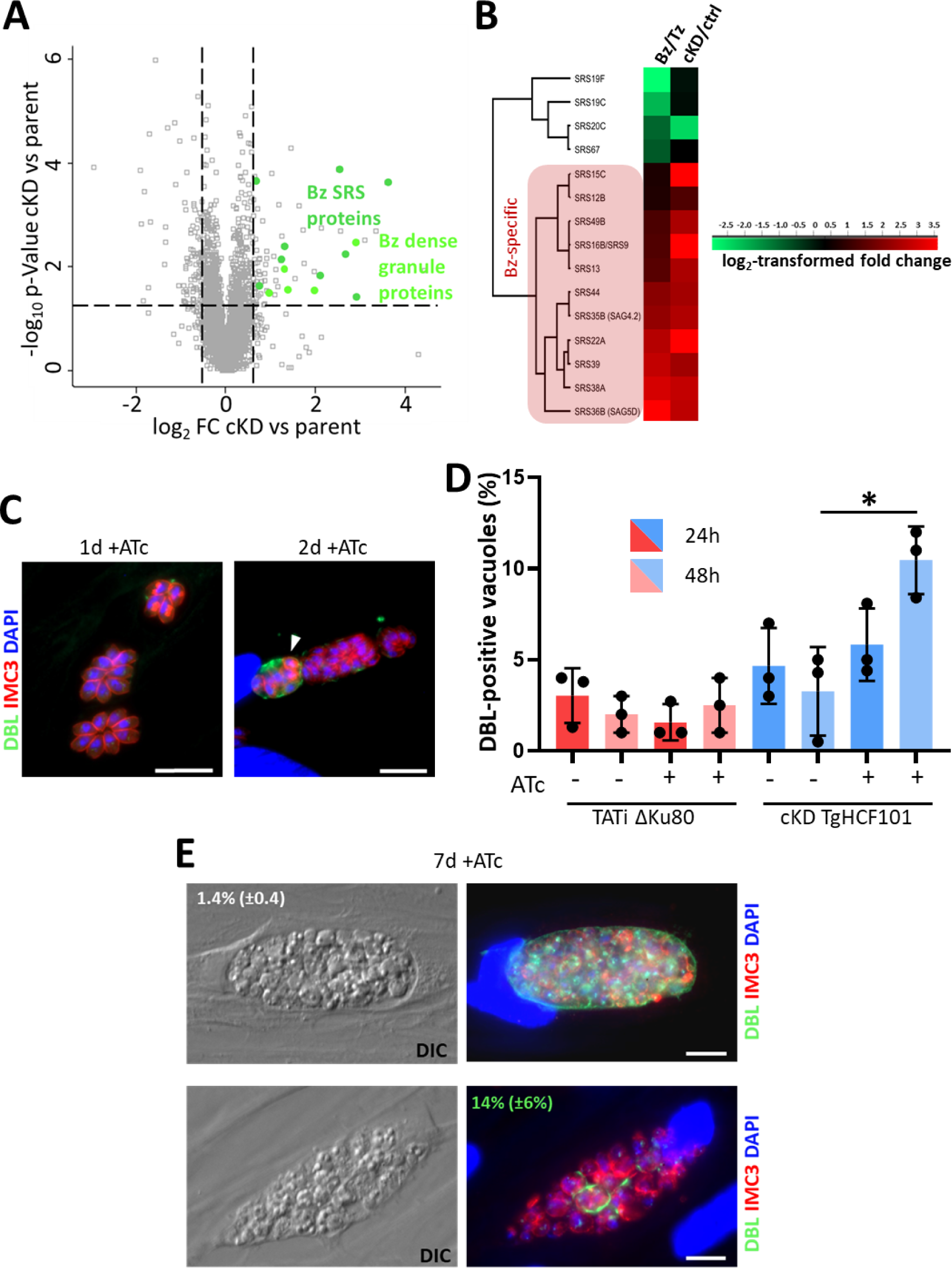
TgHCF101-depleted parasites express bradyzoite-specific markers but are unable to complete their conversion. **A.** Volcano plot showing differential expression of proteins impacted by TgHCF101 depletion after 72h of ATc treatment analyzed by label free quantitative proteomic. X-axis corresponds to the log_2_ of the fold-change (FC) and the Y-axis corresponds to the -log_10_ of the p-value, when comparing cKD-TgHCF101 expression values to the TATi ΔKu80 parental cell line. Statistical analyses were performed with ANOVA on 4 independent biological replicates. Cut-offs were set at ≤1.5- or ≥1.5-FC and p-value ≤0.05. Significant hits corresponding to stage-specific protein are highlighted in green on the graph. **B.** Clustering of bradyzoite (Bz) or tachyzoite (Tz)-specific proteins of the SRS family shows specific enrichment of bradyzoite proteins upon TgHCF101 depletion. **C.** Immunofluorescence assay of cKD-TgHCF101 treated for 24h or 48h with ATc, cyst wall is labeled with DBL, parasites periphery is outlined with anti-IMC3 antibody and DNA is stained with DAPI. Scale bar= 10µm **D.** Corresponds to the quantification of the percentage of vacuoles presenting a DBL positive signal as shown in (C). Values are represented as the mean ± SD of n=3 independent biological replicates, * p-value ≤ 0.05, Student’s *t*-test. **E.** Immunofluorescence assay of cKD-TgHCF101 treated for 7 days in the presence of ATc, cyst wall is labeled with DBL, parasites are outlined with anti-IMC3 antibody and DNA is stained with DAPI. The percentage of DBL positive vacuoles of corresponding size (more than 2 parasites per vacuole, top; or 2 parasites per vacuole or less, bottom) is specified as mean ± SD of n=3 independent biological replicates. DIC= Differential interference contrast. Scale bar= 10 µm.

These data suggest that depletion of TgHCF101 leads to a cellular stress that initiates stage conversion into the bradyzoite stage, but that the differentiation process into mature cysts cannot be completed.

### Depletion of TgHCF101 affects specifically cytosolic and nuclear Fe-S proteins

The label-free quantitative proteomic analyses also highlighted proteins which were less expressed in absence of TgHCF101. Among them, were a surprisingly large number of rhoptry bulb proteins (S3 Table, S8A Fig.). Rhoptries are club-shaped organelles that comprise a narrow tubular neck opening at the anterior pole of the parasite, and a most posterior bulbous part; the proteins secreted from these different sub-compartments are either involved in invasion and parasitophorous vacuole formation, or in the modulation of host cell defenses, respectively [47]. We assessed if there was any major impact of TgHCF101 depletion on rhoptry morphology or function. We performed IFA with anti-armadillo repeats only protein (TgARO), a protein homogenously anchored to the surface of the rhoptry membrane [48], and found no particular defect in rhoptry morphology or positioning (S8B Fig.), in accordance with our initial electron microscopy analysis (Fig. 2F). Quantification of intracellular evacuoles (rhoptry-secreted vesicular clusters) did not indicate any particular problem in the secretory capacity of the organelles upon TgHCF101 depletion (S8C Fig.). Finally, we assessed by immunoblot the expression profile of one of the potentially less-expressed rhoptry bulb protein but found only a slight alteration in the expression profile (a modest increase in the non-mature from of the protein), but not in the overall amount of protein (S8D Fig.). We conclude that although TgHCF101 depletion may lead to a secondary impact on rhoptry content, it does not extensively impact organelle morphology or secretory function.

Given the potential implication of TgHCF101 in the biogenesis of Fe-S proteins, we also specifically searched for putative Fe-S proteins in the less expressed proteins highlighted by the label-free quantitative proteomic analysis. We have previously estimated the Fe-S proteome using a computational predicting metal-binding sites in protein sequences [49], which, coupled with the data from global mapping of *T. gondii* proteins subcellular location by HyperLOPIT spatial proteomics [50], allows predicting client proteins present in the mitochondrion, apicoplast or cytosol [19]. We found three Fe-S proteins that were less expressed upon TgHCF101 depletion (Fig. 5A). Interestingly, they were all connected to the CIA pathway as they were NAR1, a Fe-S carrier to the CIA core machinery [15], and two Fe-S proteins depending on the CIA machinery for their maturation [51]: ABCE1 (Rli1 in yeast), a cytosolic ribosomal recycling factor and NTHL1, a nuclear DNA base-excision repair enzyme. In order to verify the proteomics data, we tagged TgABCE1 with a myc epitope in the context of the cKD HA-TgHCF101 mutant (S9A and B Fig.). IFA showed a decrease in the cytosolic TgABCE1 signal (Fig. 5B), which was supported by quantitative immunoblot analysis indicating a statistically significant decrease in TgABCE1 upon treatment of cKD HA-TgHCF101 mutant with ATc (Fig. 5C, D).

We did not manage to tag TgNTHL1 to verify if its abundance was effectively affected by TgHCF101 depletion. The catalytic subunit of DNA polymerase delta (TgPOLD1), which is another nucleus-associated Fe-S protein, was found to be decreasing in abundance in the CIA-related TgABCB7L mutant [40]. Our quantitative proteomic analysis did not highlight a particular decrease in TgPOLD1 abundance upon TgHCF101 depletion: it was found to be stable, or even slightly more abundant (by a 1.4-fold), after two days of TgHCF101 depletion. Yet, we knew it could be tagged and decided to use this to investigate more thoroughly a potential global impact TgHCF101 depletion on nuclear and cytoplasmic Fe-S proteins. We thus created a conditional knockdown line of TgHCF101 in the POLD1-HA background [40] and verified that it displayed a similar phenotype to the original cKD HA-TgHCF101 cell line (S10 Fig.). In the TgABCB7L mutant, TgPOLD1 abundance was found to be sharply decreasing early, before any noticeable impact on TgABCE1 [40]. In contrast, upon TgHCF101 depletion we observed a limited and delayed decrease in TgPOLD1 abundance (Fig. 6E, F, G) and intriguingly, TgPOLD1 levels even seemed to increase in the first two days of TgHCF101 depletion (Fig. 6F, G). This is in sharp contrast with the more rapid and complete impact on TgABCE1 (Fig. 6B, C, D). Of note, IFAs highlighted that although TgHCF101 depletion did not lead to a marked loss of TgPOLD1, in some parasites the protein remained in the cytoplasm instead of being translocated to the nucleus (Fig. 6E), which in fact corresponds to a stage where we have shown that DNA content of the parasite is affected (Fig. 2G).

**Figure 6.**
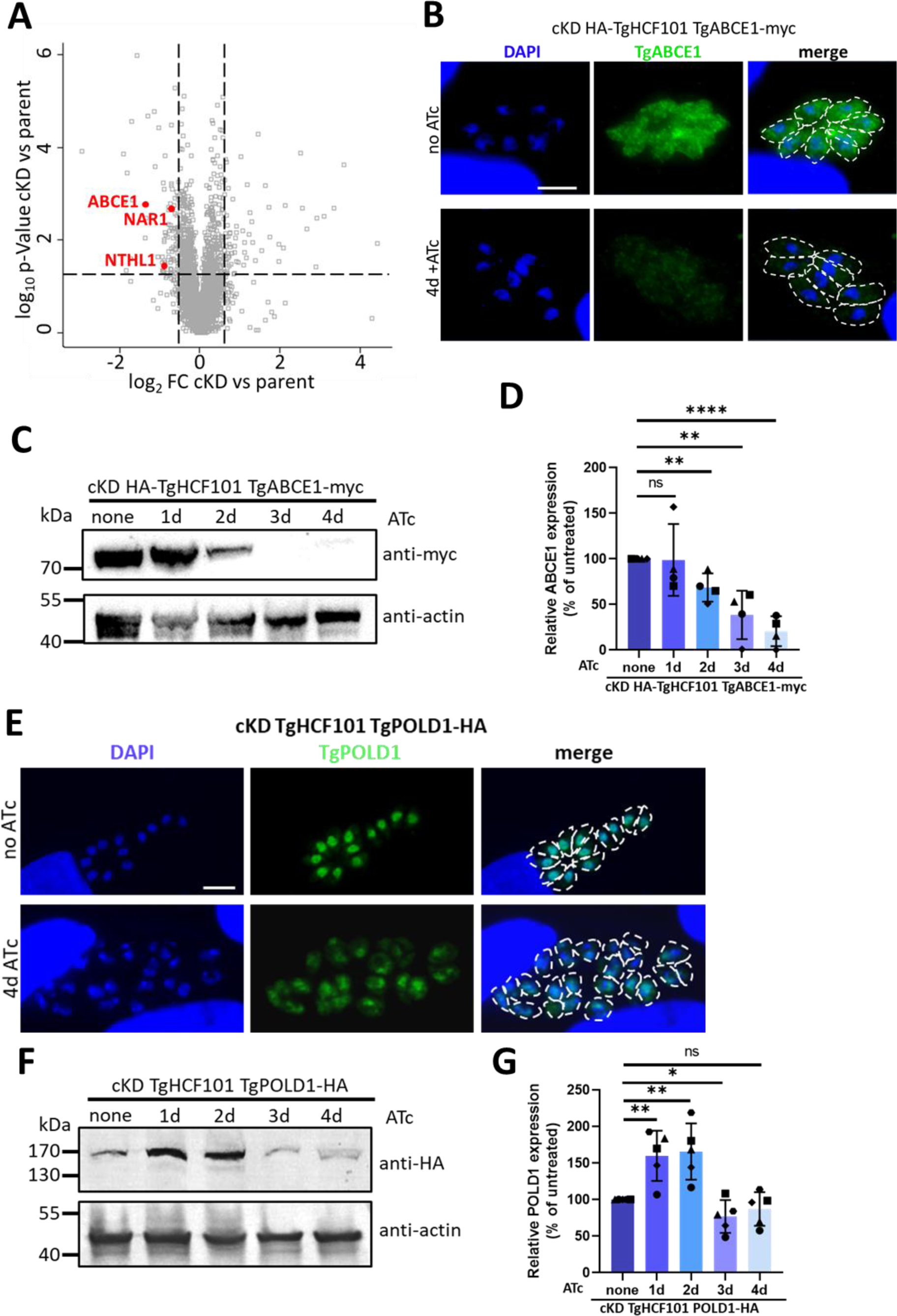
TgHCF101 is involved in Fe-S cluster biogenesis of the CIA pathway. **A.** Volcano plot showing differential expression of proteins impacted by TgHCF101 depletion after 72h of ATc treatment analyzed by label free quantitative proteomic. Cut-offs were set at ≤1.5- or ≥1.5-fold change (FC) and p-value ≤0.05. Significant hits corresponding to predicted Fe-S cluster proteins are highlighted in red on the graph. **B.** Immunofluorescence assay for the detection of myc-tagged TgABCE1 in the cKD TgHCF101 genetic background. Parasites were pre-incubated for 48h in the presence of ATc and allowed to invade HFF coated coverslips for another 48h in the presence of ATc. The control (no ATc) was infected 24h prior to fixation. TgABCE1 was detected with an anti-myc antibody and DNA was stained with DAPI. Parasites periphery is outlined by white dotted lines. Scale bar=5 µm. **C.** Immunoblot analysis of TgABCE1 abundance shows decrease upon TgHCF101 depletion following up to 4 days of treatment with ATc of the cKD HA-TgHCF101 TgABCE1-myc cell line. TgABCE1 was detected with anti-myc antibody and anti-actin antibody was used as a loading control. **D.** Decrease of TgABCE1 expression upon TgHCF101 depletion was quantified by band densitometry analysis and normalized on the loading control of each respective lane. The relative abundance of TgABCE1 is presented as a percentage relative to the untreated control, set as 100%, for each biological replicate. Values are represented as the mean ± SD from n=4 independent biological replicates, ** p-value ≤ 0.01, **** p-value ≤ 0.0001; ns, not significant (p-value ≥ 0.05), Student’s *t*-test. **E.** Immunofluorescence assay for the detection of HA-tagged TgPOLD1 in the cKD TgHCF101 genetic background. Parasites were pre-incubated for up to four days in presence of ATc on HFF-coated coverslips. The control (no ATc) was infected 24h prior to fixation. TgPOLD1 was detected with an anti-HA antibody and DNA was stained with DAPI. Parasites periphery is outlined by white dotted lines. Scale bar=5 µm. **F.** Immunoblot analysis of TgPOLD1 abundance in conditions of TgHCF101 depletion following up to 4 days of treatment with ATc of the cKD TgHCF101 TgPOLD1-HA cell line. TgPOLD1 expression was detected with anti-HA antibody and anti-actin antibody was used as a loading control. **G.** Changes in TgPOLD1 expression upon TgHCF101 depletion was quantified by band densitometry analysis and normalized on the loading control of each respective lane. The relative abundance of TgPOLD1 is presented as a percentage relative to the untreated control, set as 100%, for each biological replicate. Values are represented as the mean ± SD from n=5 independent biological replicates, * p-value ≤ 0.05; ** p-value ≤ 0.01; ns, not significant (p-value ≥ 0.05), Student’s *t*-test.

These results indicate that while perturbing TgHCF101 expression potentially has consequences on several proteins associated with the CIA machinery, it may be more specifically important for the maturation of the TgABCE1 cytosolic Fe-S protein.

### TgHCF101 is likely a Fe-S transfer protein of the CIA complex

To get further insights on the role played by TgHCF101 in the CIA machinery, we next performed co-immunoprecipitations (co-IPs) and mass spectrometry identification of associated proteins, comparing lysates of cKD HA-TgHCF101 parasites grown in the presence of ATc or not (S4 Table). Strikingly, we pulled-down all three *T. gondii* homologues of the core components of the CTC: MET18, CIA1 and AE7 (a MIP18 family protein) (Fig. 7A). We managed to tag the TgCIA1 protein with a myc epitope tag in the context of the HA-tagged TgHCF101 cell line (S9C and D Fig.) and performed a reverse IP by which we managed to specifically pull-down TgHCF101, confirming their interaction (Fig. 7B). It should be noted that this result was only obtained when this experiment was performed after chemical crosslinking, suggesting a possibly indirect or transient interaction between the proteins.

**Figure 7.**
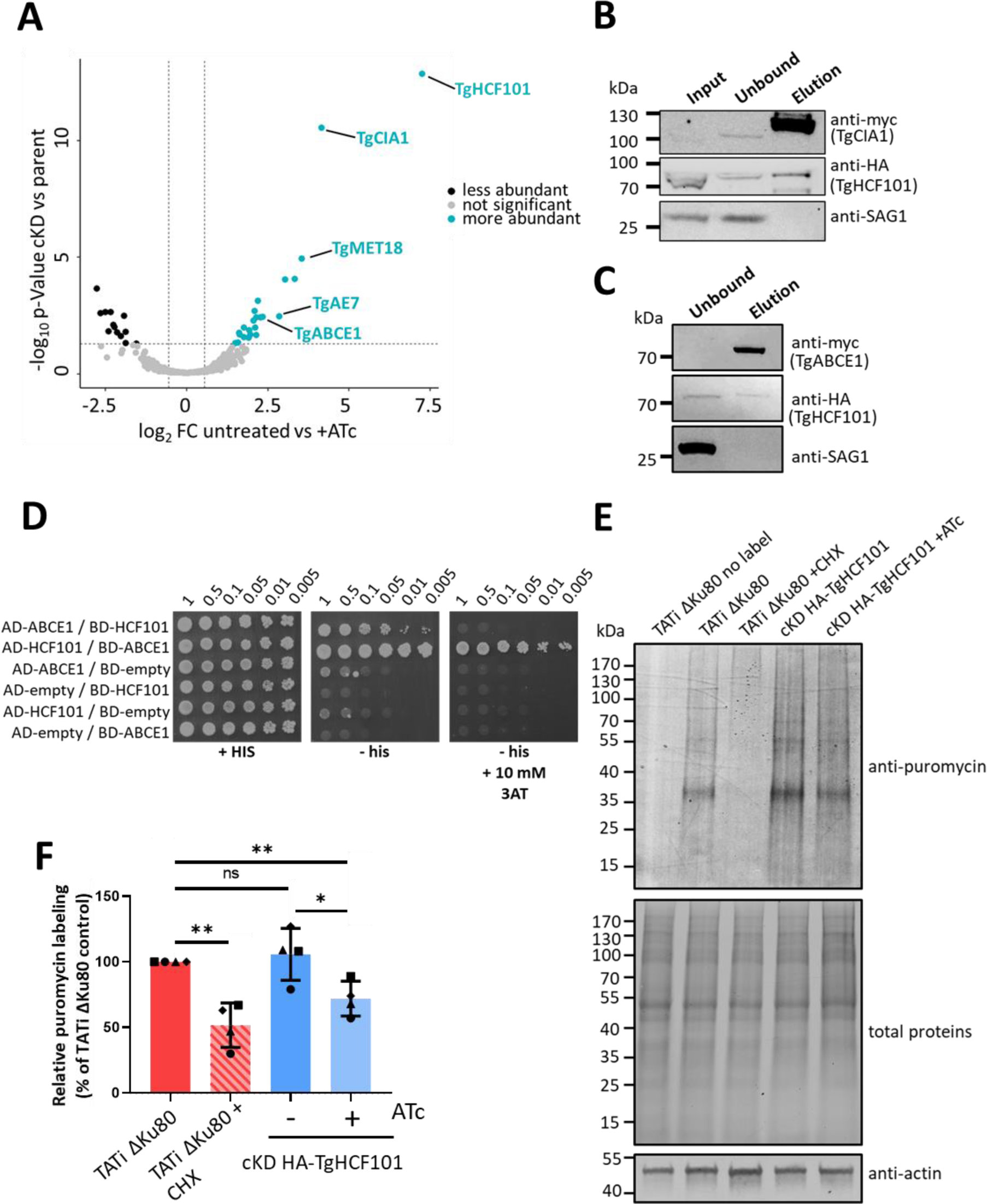
TgHCF101 is associated to the CIA targeting complex and specifically interacts with TgABCE1. **A.** Volcano plot showing differential expression of TgHCF101 and co-immunoprecipitated proteins in the cKD HA-TgHCF101 cell line after TgHCF101 depletion (with 72h of ATc treatment) or not, as analyzed by quantitative proteomic. Cut-offs were set at ≤1.5- or ≥1.5-fold change and p-value ≤0.05. Upregulated proteins are highlighted in blue, significant hits corresponding to predicted proteins of the CIA targeting complex and target protein TgABCE1 were annotated on the graph. **B.** Immunoblot analysis of a reverse co-immunoprecipitation assay of the myc-tagged TgCIA1 protein in the cKD HA-TgHCF101 background shows specific co-immunoprecipitation of TgHCF101, which is absent upon depletion by ATc. The anti-SAG1 antibody was used as a control for unspecifically bound proteins. **C.** Immunoblot analysis of a reverse co-immunoprecipitation assay of the myc-tagged TgABCE1 protein showing TgHCF101 is co-immunoprecipitating. The anti-SAG1 antibody was used as a control for unspecifically bound proteins. **D.** TgHCF101 interacts with TgABCE1 in a Gal4-based yeast two-hybrid assay. YRG2 cells co-transformed with AD-and BD-fusion proteins were grown to stationary phase, then serially diluted to OD_600_ values ranging from 1 to 5 x 10⁻³ before being spotted onto a control plate (+His, upper panel) to assess cell viability, and a Y2H test plate (-His, middle panel) to assess interaction. Plates were incubated at 30°C, and yeast growth was recorded after 5 days. As shown in the middle panel, TgHCF101 exhibited a strong interaction with TgABCE1, independent of cloning orientation. The strongest interaction was observed between AD-TgHCF101 and BD-TgABCE1, with co-transformed cells growing at the lowest dilution tested in the presence of 10 mM 3-aminotriazole (3AT) as a competitive inhibitor. Neither TgHCF101 nor TgABCE1 alone showed HIS3 transactivation in the presence of 3AT. These results are representative of 3 independent experiments. **E.** Immunoblot analysis of puromycin incorporation in the parental (TATi ΔKu80) and cKD TgHCF101 cell lines untreated or treated with ATc for 72h. TATi ΔKu80 treated with cycloheximide (CHX) was used as a control for translation inhibition. The puromycin signal was detected with anti-puromycin antibody, and total protein content was visualized by stain-Free imaging technology. Anti-actin antibody was also used as a loading control. **F.** Variation of puromycin incorporation in the different conditions was quantified by band densitometry and normalized on the total protein content of each respective lane. Puromycin labeling is presented as a percentage relative to the untreated control, set as 100% for each biological replicate. Values are represented as the mean and SD of 4 independent biological replicates; * p-value ≤ 0.05; ** p-value ≤ 0.01; ns, not significant (p-value ≥ 0.05), Student’s *t*-test.

Another interesting candidate that was found co-immunoprecipitated with TgHCF101 was TgABCE1, strengthening the possibility of an important relationship between the two proteins. We next performed a reverse IP and could recover small amounts of TgHCF101 co-eluting specifically with TgABCE1 (Fig. 7C). To further investigate this potential interaction, we performed yeast two-hybrid assay, based on interaction-dependent transactivation of the *HIS3* reporter gene (Fig. 7D). This experiment confirmed that there is a direct interaction between TgHCF101 and TgABCE1, and highlighted a particularly strong interaction, as it was retained when using 3-aminotriazole, a competitive inhibitor of the *HIS3* gene product (Fig. 7D). While this confirms that TgABCE1 is likely a client protein of TgHCF101, we could not confirm a direct interaction between TgHCF101 and TgNTHL1 using the same method (S11 Fig.). Given the known implication of ABCE1 in translation, we next evaluated the impact of TgHCF101 depletion on translation by using a puromycin-based assay (Fig. 7E). Puromycin can mimic aminoacyl-tRNA and becomes covalently attached to nascent peptides, thereby allowing the evaluation of protein elongation rates with an anti-puromycin antibody [52]. Using this method, we could show that TgHCF101 depletion leads to a modest but a statistically significant decrease in the overall translation rate (Fig. 7E, F).

Together, these results suggest that TgHCF101 acts as a Fe-S transfer protein to the translation regulator TgABCE1.

## Discussion

Fe-S clusters are universal among living organisms, where they play key roles in many important biological processes as cofactors of proteins involved for instance in housekeeping functions like respiration, photosynthesis, as well as genome expression or maintenance [6]. These cofactors were acquired early during evolution, and thus they played a fundamental role in the evolution of the eukaryotic cells as they were inherited through endosymbiosis to organelles such as the mitochondrion or plastids [5]. Different Fe-S cluster synthesis pathways show globally-conserved mechanistic and biochemical features to assemble and transfer the clusters prior to transferring them to client proteins [7]. However, beyond a core Fe-S synthesizing machinery, given the large diversity of eukaryotic lineages, it is likely that some have evolved some specific features.

For instance proteins of the MRP/NBP35 family (NBP35, Cfd1, Ind1 and HCF101), have in common a central P-loop NTPase domain [27], but are either involved in scaffolding or transfer of Fe-S clusters at specific subcellular locations like the mitochondrion, the plastid, or in the cytosol [12,25,26,53]. Presence of MRP family members in bacteria that are able to bind Fe-S clusters point to ancient and conserved function linked to Fe-S biogenesis for these proteins [54]. Another feature shared by members of this family is their ability to form homo- or hetero-dimers bridging one Fe-S cluster at their interface through two conserved and functionally essential Cys residues [55]. Interestingly, HCF101 also presents at its N-terminal a MIP18-like domain that constitutes most of a bacterial protein called SufT that is involved in Fe-S maturation [29,54,56], as well as a C-terminal domain whose function is not yet defined (DUF971). The presence of additional domain besides the conserved NTPase core domain in members of the MRP family like HCF101 suggests that they may have additional or more specialized functions. Another clue of evolutionary specialization comes from the different localization of protein isoforms in different species, hinting for a possible association to different Fe-S cluster biogenesis machineries. A recent study suggested a wide diversity in localization for HCF101 paralogs across eukaryotes, with isoforms potentially localizing to the mitochondrion in *Tetrahymena* or in the cytosol in *T. gondii* [24]. More precisely, in some chromalveolates a mitochondrial HCF101 isoform likely replaced Ind1 for the maturation of Fe-S subunits of the complex I of the mitochondrial electron transport chain. Yet, as *T. gondii* lacks complex I, it has no Ind1 homolog and no particular need for a mitochondrion-located HCF101 isoform either. Moreover, while HCF101 was initially characterized as a chloroplast-associated protein with a specialized function in maintaining PSI homeostasis, our study confirms that although *T. gondii* harbors a plastid, TgHCF101 is expressed in the cytosol (Fig. 1F, S2A Fig.), which is consistent with the absence of photosynthesis-related pathways in *T. gondii*. Our study establishes for the first time that the cytosolic expression of TgHCF101 is functionally related to the CIA pathway (Fig. 8).

**Figure 8.**
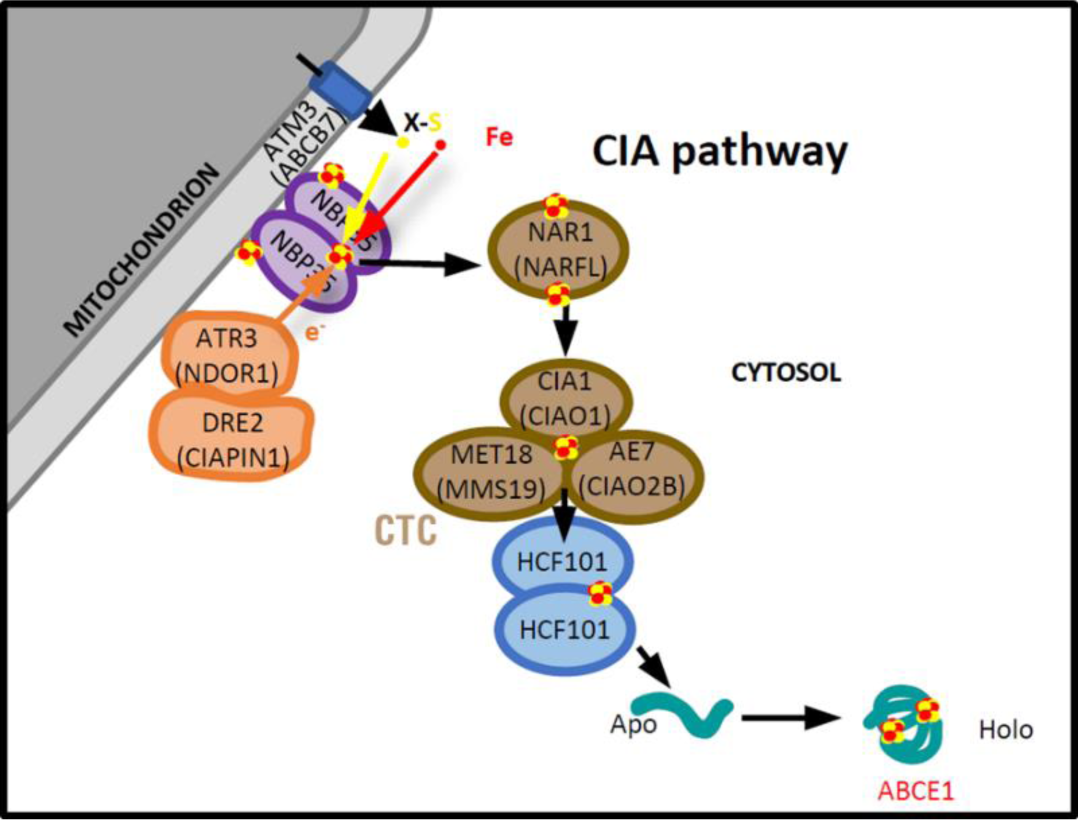
Schematic representation of the putative organization of the CIA pathway in *T. gondii.* This scheme places TgHCF101 as a Fe-S transfer protein from the CIA targeting complex (CTC) to client protein ABCE1. Plant nomenclature was used for the purpose of the figure, but names for human homologs are mentioned between brackets when appropriate.

Previous functional investigations of the CIA pathway in *T. gondii* focused on TgABCB7L, a mitochondrial transporter presumably involved in providing a sulfur-containing precursor for further processing by the cytosolic Fe-S assembly machinery [40] and on TgNBP35, a Fe-S scaffolding protein of the CIA that displays an unusual association with the outer mitochondrial membrane in the parasites [20]. The mitochondrial localization of these proteins, which are quite upstream in the CIA machinery, suggests that the assembly of cytosolic Fe-S clusters likely happens at the cytosolic face of the mitochondrion in *T. gondii*, and then probably shuttle through the *T. gondii* NAR1 homologue to the CTC for subsequent transfer to client proteins (Fig. 8). CTC proteins have so far not been functionally investigated in the parasite, but both TgABCB7L and TgNBP35 are essential for parasite growth and were both shown to be important for TgABCE1 stability [20,40]. The TgNAR1 mutant that we generated in this study also shows that this CIA shuttle protein is essential for parasite fitness (S7D Fig.). Combining genome-wide data for potential essentiality and localization [50,57] to metal-binding site prediction algorithms [49], our previous assessment of the putative *T. gondii* cytosolic and nuclear Fe-S proteome highlighted proteins involved in key functions like genome reparation and maintenance, mRNA synthesis and protein expression [19]. It is thus unsurprising to see that *T. gondii* mutants of the CIA pathway are severely impaired in growth. Our quantitative proteomics analysis of the TgHCF101 mutant did not reveal an impact on a large number of nuclear or cytosolic Fe-S proteins, which may indicate that this protein could be involved in cluster transfer to a reduced subset of Fe-S proteins. Besides the decrease in abundance of TgNAR1, strengthening the evidence of TgHCF101 involvement in the CIA pathway, quantitative proteomics data revealed a decrease in two potential client Fe-S proteins: TgABCE1 and TgNTHL1. While we could establish a direct association of TgHCF101 with TgABCE1 by co-immunoprecipitations and yeast two-hybrid (Fig. 7A, C, D), we did not manage to tag TgNTHL1 to perform co-immunoprecipitations and we did not obtain proof of a direct TgHCF101/TgNTHL1 interaction by yeast two-hybrid. Besides, when we tested the impact of TgHCF101 depletion on TgPOLD1, a Fe-S protein known to be affected by the disruption of the CIA pathway [40], we saw a limited and delayed effect compared with the one we observed on TgABCE1 (Fig. 6). Whether or not TgHCF101 is able to transfer clusters to Fe-S proteins besides TgABCE1 is still an open question, yet, so far, the evidence we gathered points primarily to TgABCE1 as its main client protein.

Our phenotypic analysis of TgHCF101-depleted parasites revealed cell division and DNA replication problems (Fig. 2D, G) that might result from an impact on Fe-S protein-mediated DNA maintenance, but may also be secondary (and more general) effects of TgHCF101 depletion. In addition, through perturbation of TgABCE1 function, we measured a decrease in the overall translation rate in the parasites (Fig. 7E, F), that could contribute to their decreased fitness and growth problems. Upon two days of TgHCF101 depletion, parasites also started to display an accumulation of LDs (Fig. 3). LDs are known to be induced as part of an integrated stress response to cellular injuries and may be induced in conditions such as oxidative stress [43]. For instance, during the particular Fe-induced cell death called ferroptosis, iron excess combined to oxygen leads to the accumulation of reactive oxygen species (ROS). This leads to subsequent peroxidation of lipids, which in turn induces the accumulation of lipid droplets that act as antioxidant organelles to control polyunsaturated fatty acid storage in triglycerides in order to reduce membrane lipid peroxidation [58]. Although it is possible that perturbation of the CIA pathway created local imbalance in cytosolic Fe concentration and ROS-dependent stress that contributed to the demise of the parasites, it is to note that our previous work has shown that general Fe deprivation has a similar LD-inducing effect on the parasites [59]. So, altogether, this may rather suggest that specific CIA-dependent Fe-S protein(s) that remain to be identified could be involved in regulating LDs. Even more so than the induction of LDs, initiation of stage conversion to the bradyzoite persistence form is a hallmark of the stress response in *T. gondii* [37]. Upon depletion of TgHCF101 we could detect early signs of differentiation (Fig. 5), but the parasites were largely unable to progress to full stage conversion and died instead (Fig. 2C, Fig. 5E). Importantly, overall our data show that TgHCF101 is essential for parasite viability, and although our findings point towards an involvement of this protein in the eukaryote-conserved CIA pathway, its specific absence from the mammalian hosts of the parasite makes it a good potential drug target. This calls for further structure/function studies of TgHCF101 in order to potentially design specific inhibitors.

There is a vast repertoire of Fe-S client proteins with a diverse range of structures and roles, and one important conundrum remaining in the Fe-S research field is resolving functional specificity of the transfer proteins during the addition of Fe-S to apoproteins. Key investigations in budding yeast allowed the identification of mutually exclusive sub-complexes of the CTC that can transfer Fe-S clusters to different proteins [51,60]. In particular, it was shown that a complex of two small proteins, Yae1 and Lto1, functions as a target-specific adaptor to recruit the yeast homologue of ABCE1 (called Rli1) to the generic CIA machinery [60]. These adaptor proteins are well-conserved in phylogenetically-close opisthokonts like humans, in which they have also been shown to be important for ABCE1 maturation [61,62]. However, Yae1 and Lto1 are clearly not conserved across all eukaryotes but found essentially in fungi, metazoan and plants (S12 Fig.), and thus noticeably they do not seem to have homologues in the Alveolata superphylum that includes apicomplexan parasites. So, while ABCE1 is arguably one of the most conserved proteins in evolution and is universally present in all eukaryotes [63], the factors driving its CIA-dependent maturation seem phylogenetically divergent. We have now established that in *T. gondii*, HCF101 is associated to the CTC and likely plays that role in this organism. Given sequence conservation of HCF101 homologs, it is also likely to be the case in other apicomplexan parasites and even possibly in one isoform present in the photosynthetic relative *C. velia* (Fig. 1A, B, S1 Fig.). Our results highlight the complex evolutionary adaptation that accompanied the maturation of eukaryotic Fe-S proteins and calls for in-depth structural analysis of this interaction. Not only this would provide invaluable evolutionary insights into the molecular machinery supporting Fe-S cluster transfer, but also may provide new prospects for interfering specifically with an essential function as a new strategy against apicomplexan-caused diseases.

## Materials and methods

### Cell culture

*Toxoplasma gondii* RH tachyzoites and derived transgenic cell lines generated in this study were routinely maintained through passages in human foreskin fibroblasts (HFFs) monolayer (ATCC CRL-1634). HFFs and parasites were cultured in standard Dulbecco’s Modified Eagle’s Medium (DMEM, supplemented with 5% decomplemented fetal bovine serum (FBS), 2 mM L-glutamine, 100 U/mL penicillin and 100 µg/mL streptomycin (Gibco) in a controlled atmosphere at 37°C and with 5% CO_2_. For immunoprecipitation assays, parasites were grown in Vero cells (ATCC, CCL-81) in the same conditions as for the HFFs.

Transgenic cell lines were generated in the TATi ΔKu80 genetic background line lacking the *Ku80* gene and expressing the TATi transactivator required for the TetOff system [64]. For positive selection of transgenic parasites bearing resistance cassettes for expressing dihydrofolate reductase thymidylate synthase (DHFR-TS) or chloramphenicol acetyltransferase (CAT) were grown with 1 µM pyrimethamine or 20 µM chloramphenicol (Sigma-Aldrich, SML3579, C0378), respectively. Conditional depletion in the TetOff conditional knockdown lines was achieved by incubation with 0,5 µg/mL anhydrotetracycline (ATc, Fluka 37919) for the indicated duration of the assay.

### Bioinformatic analyses

Genomic and protein sequences were retrieved from http://www.toxodb.org or https://www.ncbi.nlm.nih.gov/protein/. Protein alignments were performed from eukaryotic MRP family proteins (S1 Table), whose sequences were obtained from Grosche et al. [65], and updated by manual curation. Iterative rounds of alignment were performed with MAFFT v.7.475 using the E-INF-i refinement method (adapted for several conserved motifs that are embedded in long unalignable regions) https://mafft.cbrc.jp/alignment/server/index.html [66]. Sequence trimming and optimization was then performed with BMGE v.1.12 (https://ngphylogeny.fr/tools/tool/273/form) [67], with a minimal block size of 1. A phylogeny was then inferred from this dataset by maximum likelihood using the IQ-TREE software v.2.1.2 (http://www.iqtree.org/) [68], with 1,000 ultrafast bootstrap replicates [69]. Phylogenetic trees were rendered using the MEGA software v.11.0.13 (https://www.megasoftware.net/) [70] or with iTOL (https://itol.embl.de/) [71].

Transit peptide and subcellular localization predictions were performed using the ChloroP 1.1 (https://services.healthtech.dtu.dk/services/ChloroP-1.1/), IPSORT (http://ipsort.hgc.jp/), and Localizer 1.0.4 (http://localizer.csiro.au/) algorithms. Domain searches were performed using the Interpro database (https://www.ebi.ac.uk/interpro/).

### Generating a GFP-tagged TgHCF101 cell line

cDNA corresponding to the *TgHCF101* gene (TGGT1_318590) was amplified by PCR with primers ML4715 and ML4716 (all primers used in the present study are listed in S5 Table) and sub-cloned using XhoI and KpnI into the pEZS-NL vector (D. Ehrhardt, https://deepgreen.dpb.carnegiescience.edu/cell%20imaging%20site%20/html/vectors.html) for a C-terminal GFP fusion. The TgHCF101-GFP cassette was them amplified by PCR using the ML4817 and ML4818 primers and cloned using BclI and EcoRV in a BglII/EcorV-digested pTUB-IMC1-TdT vector [72] to drive the expression from a tubulin promoter. Tachyzoites were transfected with 100 µg of plasmid and observed by fluorescence microsocopy.

### Generating a conditional TgHCF101 knock-down cell line

To generate the construct for the tetracycline-regulated conditional depletion of TgHCF101 and add a N-terminal HA tag, we digested the DHFR-TetO7Sag4 plasmid [72] by BglII and inserted a fragment coding for a single HA tag generated through hybridization of the ML4924 and ML4925 oligonucleotides. Then, a 1,066 bp fragment corresponding to the 5’ coding part of TgHCF101 was amplified by PCR with primers ML4926 and ML4927 and inserted after digesting with BglII and NotI, to yield the DHFR-TetO7Sag4-HA-TgHCF101 plasmid. The TATi ΔKu80 cell line was transfected with 80 µg of the BsiWI-linearized plasmid. Transgenic parasites, named cKD HA-TgHCF101, were selected with pyrimethamine and cloned by limiting dilution. Positive clones were verified by PCR with primers ML1771 and ML6043.

### Generating complemented cell lines

The cKD TgHCF101-HA cell line was complemented by adding an extra copy of the *TgHCF101* gene under the control of a tubulin promoter at the *UPRT* locus. The entire *TgHCF101* cDNA sequence (1935 bp) was amplified by PCR using primers ML6409/ML6410 from the plasmid pGBKT7-myc-HCF101 to express a N-terminal myc-tagged HCF101 copy. This sequence was then cloned downstream of the tubulin promoter in the pUPRT-TUB vector [64], resulting in the pUPRT-myc-TgHCF101 plasmid. The plasmid was subsequently linearized prior to transfecting the mutant cell line, together with a plasmid expressing Cas9 and a *UPRT*-specific guide RNA under the control of a U6 promoter [73]. Transgenic parasites were selected using 5 µM 5-fluorodeoxyuridine (Sigma-Aldrich) and cloned by serial limiting dilution to obtain the cKD TgHCF101 WT complemented cell line.

For complementation with the copy of TgHCF101 lacking the LEW tripeptide motif, the cDNA sequence was amplified by PCR using primers ML6409/ML6412 from the plasmid pGADT7-myc-HCF101, to remove the tripeptide motif in the C-terminal region. All constructs were verified by sequencing. Cloning and transfection were performed as previously described to establish the cKD TgHCF101 -LEW comp cell line. For both cell lines correct integration at the *UPRT* locus was verified by PCR using primers ML6044 and ML6396.

### Generating a conditional TgNAR1 knock-down cell line

We digested the DHFR-TetO7Sag4 plasmid by BglII and then, a fragment corresponding to the 5’ coding part of *TgNAR1* was amplified by PCR with primers ML5862 and ML5863 and inserted after digesting with BamHI and NotI to yield the DHFR-TetO7Sag4-TgNAR1 plasmid. The TATi ΔKu80 cell line was transfected with 80 µg of the NsiI-linearized plasmid. Transgenic parasites, named cKD HA-TgNAR1, were selected with pyrimethamine and cloned by limiting dilution. Positive clones were verified by PCR with primers ML1771 and ML5947.

### Tagging of ABCE1 and CIA1 in the TgHCF101 knock-down background

A CRISPR-based strategy was used to C-terminally tag proteins of interest in the cKD HA-TgHCF101 background. Guide RNAs (gRNAs) targeting the C-terminal end of the gene of interest were selected using CHOP-CHOP tool (https://chopchop.cbu.uib.no/). gRNAs were cloned into the pU6-Cas9 Universal Plasmid (Addgene, 52694) using BsaI. Donor DNA was amplified by PCR using the high fidelity KOD DNA polymerase (Novagen) to amplify a fragment containing the tag and the resistance cassette, using the pLIC-myc-CAT plasmid as a template, adding at both ends 30-nucleotide overhangs homologous to the C-terminal region of the gene of interest for homologous recombination. *TgABCE1* (TGGT1_216790) tagging was performed by using primers ML5126 and ML5127 for the gRNA and ML5128 and ML5129 to amplify donor DNA, yielding the cKD HA-TgHCF101 TgABCE1-myc cell line. Integration of the construct was controlled by PCR using GoTaq DNA polymerase (Promega) with primers ML6115 and ML4310. *TgCIA1* (TGGT1_313280) tagging was performed by using primers ML5932 and ML5933 for the gRNA and ML5934 and ML5935 to amplify donor DNA, yielding the cKD HA-TgHCF101 TgCIA1-myc cell line. Integration of the construct was controlled by PCR using GoTaq DNA polymerase (Promega) with primers ML5936 and ML4310.

### Generating a TgHCF101 knock-down cell line with HA-tagged POLD1

The POLD1-HA cell line [40] was used as a background to generate a tetracycline-regulated conditional TgHCF101 mutant. Primers ML4926 and ML4927 were used to amplify a a 1,066 bp 5’ fragment of the *TgHCF101* coding sequence and cloned using BglII/NotI in the DHFR-TetO7Sag4 plasmid [72] to yield the DHFR-TetO7Sag4-TgHCF101 plasmid. The POLD1-HA cell line was transfected with 80 µg of this plasmid linearized by NsiI. Transgenic parasites were selected with pyrimethamine and cloned by limiting dilution. Positive clones were verified by PCR with primers ML2456 and ML6043.

### Semi-quantitative RT-PCR

Semi-quantitative RT-PCR was performed as described previously [74]. Briefly, 1µg of RNA was used as a template for each RT-PCR reaction and specific primers for *TgHCF101* (ML5112/ML5113), *TgNAR1* (ML6467/ML6468) or *TUB2* (Tubulin β chain) (ML841/ML842) were used. Twenty-four cycles of PCR were performed.

### Immunofluorescence assays

For immunofluorescence assays (IFA), coverslips seeded with HFFs were infected by *T. gondii* tachyzoites. Intracellular parasites and host cell monolayer were then fixed with 4% (w/v) paraformaldehyde (PFA, diluted in phosphate-buffered saline -PBS-) for 20 minutes. After fixation, cells were permeabilized with 0.3% (v/v) Triton X-100 (diluted in PBS) for 10 minutes. Coverslips were blocked with 2% (w/v) bovine serum albumin (BSA) for 1h prior to immunolabeling with primary antibody for 1 hour. After 3 washes in PBS, corresponding secondary antibody was incubated for 1 hour. Coverslips were finally incubated with 1 µg/mL 4,6-diamidino-2-phenylindole (DAPI) for 5 minutes before 3 washes in PBS and, lastly, mounted using Immu-Mount (ThermoFisher) onto microscope slides. Primary antibodies used were prepared in 2% BSA (diluted in PBS) and used at the following concentrations: rat monoclonal anti-HA (1:1000, 3F10 Roche), mouse monoclonal anti-myc (1:100, 9E10 Sigma), rabbit anti-IMC3 (1:1000) [75], mouse anti-SAG1 (T41E5, 1:1000) [76], rabbit anti-armadillo repeats only (ARO, 1:1000) [48], rabbit anti-PDH-E2 (1:500) [22], mouse anti-F1β ATPAse (1:1000, gift of P. Bradley). Cysts were stained with biotin labeled *Dolichos Biflorus* lectin (1:300, Sigma L-6533) and detected with FITC-conjugated streptavidin (1:300, Invitrogen, SNN1008). Lipid droplets were detected with Nile Red (1 µg/mL, Sigma 72485), using the CY3/DsRed BP550/25 FT570 BP605/70 filter cube (Zeiss) on an epifluorescence microscope.

All images were acquired at the Montpellier Ressources Imagerie (MRI) facility. Observations were performed with Zeiss AxioImager Z1 and Z2 epifluorescence microscopes equipped with a Zeiss Axiocam MRm CCD camera and 63X/1.4 or 100X/1.4 Oil Plan Achromat objective. Images were processed on Zen Blue v3.6 (Blue edition) software (Zeiss). Z-stack acquisitions were processed by maximum intensity orthogonal projection when assessing lipid droplets number and area. Adjustments of brightness and contrast were applied uniformly and paired images were acquired with the same exposure time.

### Plaque and replication assays

The lytic cycle of tachyzoites was assessed by plaque assay as described previously [59]. Briefly, tachyzoites from the cKD HA-TgHCF101 transgenic cell line or parental strain (TATi ΔKu80) were allowed to invade monolayers of HFFs in the presence or absence of ATc. Parasites were cultivated for 7 days at 37°C and 5% CO_2_ and fixed with 4% (w/v) PFA (diluted in PBS) for 20 minutes. Cells were stained with 0.1% crystal violet solution (V5265, Sigma-Aldrich), washed and air-dried before imaging on an Olympus MVX10 microscope. For the reversibility assay, drug washout was carefully performed after 7 days of ATc pre-treatment and cultures were kept for another 7 days of growth, non-treated control conditions were also infected at this timepoint.

For replication assay, parasites were pre-treated in flasks for 48 hours with ATc and extracellular parasites were allowed to invade HFF monolayer on coverslips for another 24 hours before being fixed and performing parasites immunodetection by IFA with mouse anti-SAG1 antibody as described before. The number of parasites per vacuole was scored. Independent experiments were conducted three times, and 200 random vacuoles were counted for each condition.

### Immunoblotting and antibodies

Protein extracts were prepared from 10^7^ extracellular parasites resuspended in Laemmli buffer at a final concentration of 10^6^ parasites/µL. Extracts were treated with benzonase to remove DNA from samples and resolved by SDS-PAGE before being transferred on nitrocellulose membrane for subsequent protein detection. Primary antibodies used for immunodetection were resuspended at their respective working concentration in 5% (w/v) milk in TNT buffer (0.1 M Tris-HCl, pH 7.6, 0.15 M NaCl, 0.05%, Tween 20). Antibodies used in this study for immunoblot detection were rat anti-HA (1:1000, Roche), mouse anti-myc (1:100, Sigma), mouse anti-SAG1 (1:50, hybridoma), mouse anti-actin (1:25, hybridoma) [77], mouse anti-ROP7 (1:1000, T43H1) [78], mouse anti-puromycin (1:1000,12D10, MABE943 Sigma) and rabbit anti-lipoic acid (1:500, ab58724 Abcam) [79].

### Puromycin labeling

Strains of interest were grown for 3 days in the presence or absence of ATc. Freshly egressed parasites were filtered on 40µm Cell Strainer (VWR, 723-2757). After filtration and counting, parasites were treated with puromycin (100 µg/mL, puromycin dihydrochloride, Sigma) for 15 minutes at 37°C and 5% CO_2_. For translation inhibition control, parasites were treated with cycloheximide (100 µg/mL, Sigma) for 10 minutes prior to puromycin incubation. After treatment, parasites were washed in DPBS (Dulbecco’s phosphate-buffered saline, Gibco) and collected by centrifugation. Pellet was resuspended in Laemmli buffer and separated on Mini-Protean TGX Stain-free gels 12% (BioRad) activated by a UV-induced 1-minute reaction to produce tryptophan residue fluorescence in order to allow for global protein quantification, following manufacturer instructions. Proteins were then transferred to nitrocellulose membrane for immunodetection using mouse anti-puromycin (1:1000,12D10, MABE943 Sigma) and mouse anti-actin (1:25, hybridoma). Total protein content is assessed by Stain-free detection and puromycin signal is detected with secondary antibody mouse coupled with alkaline phosphatase. Both signals were quantified by densitometry using the ImageJ software.

### Electron microscopy

Parasites were pretreated with ATc for 48h and allowed to reinvade for 24h in the presence of ATc. Untreated parasites were used as a control for normal morphology. Cells were then fixed with 2.5% glutaraldehyde in cacodylate buffer 0.1 M pH7.4. Coverslips were subsequently processed using a Pelco Biowave pro+ (Ted Pella). Samples were postfixed in 1% OsO4 and 2% uranyl acetate, dehydrated in acetonitrile series and embedded in Epon 118 using the following parameters: Glutaraldehyde (150 W ON/OFF/ON 1-min cycles); two buffer washes (40 s 150 W); OsO4 (150 W ON/OFF/ON/OFF/ON 1-min cycles); two water washes (40 s 150 W); uranyl acetate (100 W ON/OFF/ON 1-min cycles); dehydration (40 s 150 W); resin infiltration (350 W 3-min cycles). Fixation and infiltration steps were performed under vacuum. Polymerization was performed at 60°C for 48 hr. Ultrathin sections at 70 nM were cut with a Leica UC7 ultramicrotome, counterstained with uranyl acetate and lead citrate and observed in a Jeol 1400+ transmission electron microscope from the MEA Montpellier Electron Microscopy Platform. All chemicals were from Electron Microscopy Sciences, and solvents were from Sigma.

### Label-free quantitative proteomics

Parasites from the TATi ΔKu80 and cKD HA-TgHCF101 cell lines were treated for 72 hours in HFF seeded in T75 cm^2^ flasks. After suitable incubation, parasites were released from host cells with a cell scraper and passed through a 25G needle before filtration on a fiber glass wool column. Parasites were pelleted and washed in Hank’s Balanced Salt Solution (HBSS, Gibco). Parasites were resuspended in lysis buffer (1% SDS, 50mM Tris HCL pH 8, 10mM EDTA pH 8) and protein quantification was determined with the bicinchoninic acid kit (Abcam). For each condition, 20 µg of protein resuspended in Laemmli buffer were resolved on a 10% SDS-PAGE for 35 minutes at 100V. Proteins were fixed with a combination of acetic acid and ethanol and stained in PageBlue Protein Staining Solution (ThermoScientific). Each lane was cut in three identical pieces which were digested with trypsin and peptide extraction was done as previously described [80].

LC-MS/MS experiments were performed using an Ultimate 3000 RSLC nano system (ThermoFisher) interfaced online with a nano easy ion source and an Exploris 240 Plus Orbitrap mass spectrometer (ThermoFisher). The .raw files were analyzed with MaxQuant version 2.0.3.0 using default settings (PMID: 19029910). The minimal peptide length was set to 6. The files were searched against the *T. gondii* proteome (March 2020, https://www.uniprot.org/proteomes/UP000005641-8450). Identified proteins were filtered according to the following criteria: at least two different trypsin peptides with at least one unique peptide, an E value below 0.01 and a protein E value smaller than 0.01 were required. Using the above criteria, the rate of false peptide sequence assignment and false protein identification were lower than 1%. Proteins were quantified by label-free method with MaxQuant software using unique and razor peptides intensities [81]. Statistical analyses were carried out using RStudio package software. The protein intensity ratio (protein intensity in mutant/protein intensity in parent) and statistical tests were applied to identify the significant differences in the protein abundance. Hits were retained if they were quantified in at least three of the four replicates in at least one experiment. Proteins with a statistically significant (p < 0.05 or 0.01 with or without Benjamini correction) quantitative ratio were considered as significantly up-regulated and down-regulated respectively. Volcano plots were generated with the Perseus software v2.0.7.0 (https://maxquant.net/perseus/). Perseus was also used for hierarchical clustering of bradyzoite-and tachyzoite-specific surface antigens of the SRS family using RNAseq data of Hehl et al. [82], available on www.Toxodb.org.

### Co-immunoprecipitation and mass spectrometry identification

Parasites of the cKD HA-TgHCF101 transgenic cell line were treated for 3 days in the presence or absence of ATc in T175 cm^2^ seeded with Vero cells. After treatment, intracellular parasites were released by scraping of the host cells and three passages through a 26G needle. To eliminate cell host debris, parasites were filtered through fiber glass wool and harvested by centrifugation at 650g for 5 min and washed three times in DPBS (Gibco). Parasites were resuspended in lysis buffer (1%NP40, 50 mM Tris-HCl pH8, 150 mM NaCl, 4 mM EDTA, supplemented with cOmplete Mini protease inhibitors mix (Roche)) and incubated overnight at 4°C on a rotating wheel. Centrifugation of insoluble material was performed at 13500g for 30 min at 4°C. The supernatant was transferred to a tube containing 50µL of anti-HA magnetic beads (ThermoFisher, 88836) for 4h at 4°C on a rotating wheel. The depleted fraction was then removed and beads were washed 5 times with lysis buffer. For the elution of immunoprecipitated proteins, beads were incubated in 50 µL of HA peptide solution at 2 mg/mL (ThermoFisher, 26184) for 1h at 37°C on a rotating wheel. The eluted proteins were mixed with Laemmli buffer and resolved on a 10% SDS-Page for 35 minutes at 100V. In-gel proteins were fixed with a combination of acetic acid and ethanol and stained with PageBlue Protein Staining Solution (ThermoScientific). Each lane was cut in three identical pieces which were digested with trypsin, peptide extraction was done as previously described [80] and mass spectrometry identification was performed as described for label-free quantitative proteomic.

Proteomic data were analyzed on R following the publicly available script of DEP analysis package (https://github.com/arnesmits/DEP, v.1.7.1). The following parameters were used for differential expression analysis: dataset was normalized by variance-stabilizing transformation and missing values were imputed via MNAR (missing not at random) method, the fold change cutoff was set at 1.5 and p-value at 0.05. Statistical analyses were performed by a differential enrichment test based on protein-wise linear models and empirical Bayes statistics [83] and p-values were adjusted by the Benjamini-Hochberg correction.

Validation of co-immunoprecipitation mass spectrometry results was performed by immunoblotting. C-terminally c-myc tagged candidates in the cKD HA-TgHC101 background were grown in the presence or absence of ATc for 48h. Parasites were harvested and subjected to co-immunoprecipitation using myc-Trap agarose beads (Chromotek, yta-20) following manufacturer’s instructions. Cross-linking of proteins was performed prior to co-immunoprecipitation for the cKD HA-TgHCF101 CIA1-myc cell line using 1% (w/v) PFA (diluted in PBS) for 5 minutes at room temperature and quenched with 125 mM ice-cold glycine. Immunoprecipitated proteins were then resolved on 10% SDS-Page prior to immunoblot analysis.

### Yeast two-hybrid

RNA was extracted from parasites using NuceloSpin RNA kit (Macherey-Nagel). cDNA was obtained using SuperScript III First Strand Synthesis SuperMix for RT-qPCR (Invitrogen) using oligodT and following the manufacturer’s instructions. Specific cDNAs were subsequently amplified using KOD DNA polymerase (Novagen). Cloning was performed with In-Fusion HD cloning kit (Takara) for each candidate into pGADT7 and pGBKT7 vectors (Clontech, Takara Bio) to allow expression of AD-(Gal4 activation domain) and BD-(Gal4 DNA binding domain) fusion proteins, respectively. Primers were designed using the InFusion Cloning Primer Design Tool (https://takarabio.com/) and are listed in S5 Table.

All experiments were performed in the Gal4-based yeast two-hybrid (Y2H) reporter strain YRG2 (MATα, *ura3-52*, *his3-200*, *ade2-101*, *lys2-801*, *leu2-3*, *112*, *trp1-901*, *gal4-542*, *gal80-538*, *lys2*::UAS_GAL1_-TATA_GAL1_-HIS3, URA3::UAS_GAL4_,*_17MERS(x3)_*-TATA_CYC1_-LacZ) (Stratagene, Agilent). Yeast cells were co-transformed by pGAD/pGBK construct pairs and selected on plates containing the minimal YNB medium (0.7% Yeast Nitrogen Base medium without amino acids, 2% glucose, 2% Difco agar) supplemented with histidine (H), adenine (A), Lysine (K), and uracil (U) according to Gietz and Woods [84]. For all Y2H interaction assays, individual co-transformed colonies were cultured on liquid YNB+HUKA, adjusted to an OD_600_ of 0.05 and dotted on YNB+UKA plates containing (+His plates) or missing (-His plates) histidine (7 µL per dot). Plates were incubated at 30°C, and cell growth was recorded after 5 days. Constructs pairs including empty vectors (AD-fusion + empty BD, or empty AD + BD-fusion) were tested first to evaluate the capacity of each protein of interest to transactivate the *HIS3* reporter gene in the absence of interactor, and to determine the appropriate 3-aminotriazole (3AT) concentration required to abolish false interactions. True positive interactions were visualized as cells growing on YNB+UKA in the absence of histidine (-his plates) and eventually in the presence of 3AT if required. Increasing 3AT concentrations were also used to challenge the strength of positive interactions. For all assays, at least three independent transformations per construct pair were performed.

### DNA content analysis

DNA content analysis by flow cytometry was performed as described previously [85]. Parasites from the cKD HA-TgHCF101 strain and from the background line TATi ΔKu80 were treated for up to four days in the presence or absence of ATc. Extracellular parasites were removed by washing with HBSS and intracellular parasites were released from host cell by scraping with a cell scraper (VWR, 734-2602), passages through 26G needles and finally filtered on a fiber glass wool column. Parasites were then fixed overnight in a solution of 70% ethanol and 30% PBS at 4°C. After fixation, parasites were washed in PBS and stained with a 30 µM propidium iodide solution for 30 min. Parasites DNA content was then analyzed by flow cytometry with an Aurora cytometer (Cytek) from the MRI facility.

### Gliding motility assay

Parasites from the parental TATi ΔKu80 and the cKD HA-TgHCF101 transgenic cell lines were grown in the presence or absence of ATc for up to 4 days. 1.10^6^ freshly egressed parasites were harvested and resuspended in 400µL motility buffer (155 mM NaCl, 3 mM KCl, 2 mM CaCl_2_, 1 mM MgCl_2_, 3 mM NaH_2_PO_4_, 10 mM HEPES, 10 mM glucose) and 100 µL of the suspension was immediately placed on poly-L-lysine coated slides and incubated at 37°C for 15 min. Unattached parasites were removed by performing three washes with PBS. Finally, attached parasites were fixed with 4% (w/v) PFA and stained with anti-SAG1 antibody. Trails length (at least 100 per condition) were captured on random fields with a 63x objective on a Zeiss AXIO Imager Z2 epifluorescence microscope and were quantified using the NeuronJ plug-in on the ImageJ software as described previously [86].

### Rhoptry secretion assay

Rhoptry secretion was quantified by performing an evacuole assay [87]. Parasites from the cKD HA-TgHCF101 strain and cKD TgARO strain were grown in the presence or absence of tetracycline for up to 72 hours. Freshly egressed parasites were then pre-treated for 10 min in DMEM supplemented with 1 µM cytochalasin D (cytD), an inhibitor of actin polymerization. Parasites were allowed to secrete their rhoptry content as they were put in contact with HFF cells for 15 min in the presence of cytD at 37°C. Parasites and HFF cells were then immediately fixed with 8% (w/v, in PBS) PFA, permeabilized with 0.1% Triton X-100 and stained with anti-ROP1 and anti-SAG1 to detect secreted rhoptry material and parasites, respectively. For each technical replicate, 20 random fields were quantified and three technical replicates were performed for each of the three biological replicates.

### Statistical analyses

Statistical analyses were generally performed with the Prism 8.3 software (Graphpad). For proteomics experiments, statistics were performed with version 4.2.1 of the R package (2022-06-23, https://www.R-project.org/) using the Differential Enrichment analysis of Proteomics Data (DEP, v.1.23.0) package. Unless specified, values are expressed as means ± standard deviation (SD).

## Data availability

All raw MS data and MaxQuant files generated have been deposited to the ProteomeXchange Consortium via the PRIDE partner repository (https://www.ebi.ac.uk/pride/archive) with the dataset identifiers PXD051549 and PXD051551.

## Acknowledgements

We thank B. Striepen, V. Carruthers, M.J. Gubbels, P. Bradley, A. MacLean and D. Soldati-Favre for the gift of antibodies or cell lines. Mass spectrometry experiments were carried out using the facilities of the Montpellier Proteomics Platform (PPM, MSPP site, BioCampus Montpellier), member of the Proteomics French Infrastructure (ProFI). We also thank the developers and the managers of the VeupathDB.org and ToxoDB.org databases, as well as scientists who contributed datasets. We acknowledge the MRI Cell Imaging Facility, member of the national infrastructure France-BioImaging for image acquisitions and cytometry analyses.

## Suppl. Table Legends

**S1 Table. Sequences used for the phylogenetic analysis displayed in Fig. 1A.** Abbreviations used in the Fig. 1A phylogenetic analysis are mentioned in red next to each sequence.

**S2 Table. Proteins with higher expression upon depletion of TgHCF101 as found by label-free quantitative proteomics.** For each protein candidate (with www.ToxoDB.org and www.Uniprot.org identifier), log_2_ of the different ratio were calculated between the mean MaxQuant LFQ values found for the cKD HA-TgHCF101 mutant and the TATi ΔKu80 parental cell line. -log_10_(pvalue) is also provided. Putative subcellular localization was obtained from the hyperLOPIT data available on ToxoDB.org, or by manual annotation. CRISPR fitness score and transcriptomic data for tachyzoites (Tz) and bradyzoites (Bz) were obtained from ToxoDB.org.

**S3 Table. Proteins with lower expression upon depletion of TgHCF101 as found by label-free quantitative proteomics.** For each protein candidate (with www.ToxoDB.org and www.Uniprot.org identifier), log_2_ of the different ratio were calculated between the mean MaxQuant LFQ values found for the cKD HA-TgHCF101 mutant and the TATi ΔKu80 parental cell line. -log_10_(pvalue) is also provided. Putative subcellular localization was obtained from the hyperLOPIT data available on ToxoDB.org, or by manual annotation. CRISPR fitness score and transcriptomic data for tachyzoites (Tz) and bradyzoites (Bz) were obtained from ToxoDB.org.

**S4 Table. Proteins identified as co-immunoprecipitating with TgHCF101 through comparative mass spectrometry analysis of immunoprecipitated extracts form the cKD HA-TgHCF101 cell line grown or not in the presence of ATc.** Proteins enriched (TgHCF101-expressing sample vs TgHCF101-depleted control) in a statistically significant way (p-value ≤ 0.05, and Benjamini-Hochberg correctin) are in green (dark green for TgHCF101 and putative iron-sulfur protein) and those less abundant are in red.

**S5 Table. Oligonucleotides used in this study.**

## References

1. Beinert H, Holm RH, Münck E. Iron-sulfur clusters: nature’s modular, multipurpose structures. Science. 1997;277: 653–659. doi:10.1126/science.277.5326.653

2. Andreini C, Putignano V, Rosato A, Banci L. The human iron-proteome. Metallomics. 2018;10: 1223–1231. doi:10.1039/c8mt00146d

3. Lill R, Freibert S-A. Mechanisms of mitochondrial iron-sulfur protein biogenesis. Annu Rev Biochem. 2020;89: 471–499. doi:10.1146/annurev-biochem-013118-111540

4. Brzóska K, Meczyńska S, Kruszewski M. Iron-sulfur cluster proteins: electron transfer and beyond. Acta Biochim Pol. 2006;53: 685–691.

5. Tsaousis AD. On the origin of Iron/Sulfur cluster biosynthesis in eukaryotes. Front Microbiol. 2019;10: 2478. doi:10.3389/fmicb.2019.02478

6. Garcia PS, D’Angelo F, Ollagnier De Choudens S, Dussouchaud M, Bouveret E, Gribaldo S, et al. An early origin of iron–sulfur cluster biosynthesis machineries before Earth oxygenation. Nat Ecol Evol. 2022;6: 1564–1572. doi:10.1038/s41559-022-01857-1

7. Braymer JJ, Freibert SA, Rakwalska-Bange M, Lill R. Mechanistic concepts of iron-sulfur protein biogenesis in Biology. Biochimica et Biophysica Acta (BBA) - Molecular Cell Research. 2021;1868: 118863. doi:10.1016/j.bbamcr.2020.118863

8. Maio N, Rouault TA. Outlining the complex pathway of mammalian Fe-S cluster biogenesis. Trends in Biochemical Sciences. 2020;45: 411–426. doi:10.1016/j.tibs.2020.02.001

9. Kispal G, Csere P, Prohl C, Lill R. The mitochondrial proteins Atm1p and Nfs1p are essential for biogenesis of cytosolic Fe/S proteins. EMBO J. 1999;18: 3981–3989. doi:10.1093/emboj/18.14.3981

10. Pondarré C, Antiochos BB, Campagna DR, Clarke SL, Greer EL, Deck KM, et al. The mitochondrial ATP-binding cassette transporter Abcb7 is essential in mice and participates in cytosolic iron– sulfur cluster biogenesis. Human Molecular Genetics. 2006;15: 953–964. doi:10.1093/hmg/ddl012

11. Zuo J, Wu Z, Li Y, Shen Z, Feng X, Zhang M, et al. Mitochondrial ABC transporter ATM3 is essential for cytosolic iron-sulfur cluster assembly. Plant Physiol. 2017;173: 2096–2109. doi:10.1104/pp.16.01760

12. Camire EJ, Grossman JD, Thole GJ, Fleischman NM, Perlstein DL. The yeast Nbp35-Cfd1 cytosolic iron-sulfur cluster scaffold is an ATPase. J Biol Chem. 2015;290: 23793–23802. doi:10.1074/jbc.M115.667022

13. Netz DJA, Stümpfig M, Doré C, Mühlenhoff U, Pierik AJ, Lill R. Tah18 transfers electrons to Dre2 in cytosolic iron-sulfur protein biogenesis. Nat Chem Biol. 2010;6: 758–765. doi:10.1038/nchembio.432

14. Kohbushi H, Nakai Y, Kikuchi S, Yabe T, Hori H, Nakai M. Arabidopsis cytosolic Nbp35 homodimer can assemble both [2Fe–2S] and [4Fe–4S] clusters in two distinct domains. Biochemical and Biophysical Research Communications. 2009;378: 810–815. doi:10.1016/j.bbrc.2008.11.138

15. Balk J, Pierik AJ, Netz DJA, Mühlenhoff U, Lill R. The hydrogenase-like Nar1p is essential for maturation of cytosolic and nuclear iron-sulphur proteins. EMBO J. 2004;23: 2105–2115. doi:10.1038/sj.emboj.7600216

16. Kassube SA, Thomä NH. Structural insights into Fe–S protein biogenesis by the CIA targeting complex. Nat Struct Mol Biol. 2020;27: 735–742. doi:10.1038/s41594-020-0454-0

17. Sanchez SG, Besteiro S. The pathogenicity and virulence of Toxoplasma gondii. Virulence. 2021;12: 3095–3114. doi:10.1080/21505594.2021.2012346

18. Sheiner L, Vaidya AB, McFadden GI. The metabolic roles of the endosymbiotic organelles of Toxoplasma and Plasmodium spp. Curr Opin Microbiol. 2013;16: 452–458. doi:10.1016/j.mib.2013.07.003

19. Pamukcu S, Cerutti A, Bordat Y, Hem S, Rofidal V, Besteiro S. Differential contribution of two organelles of endosymbiotic origin to iron-sulfur cluster synthesis and overall fitness in Toxoplasma. Soldati-Favre D, editor. PLoS Pathog. 2021;17: e1010096. doi:10.1371/journal.ppat.1010096

20. Aw YTV, Seidi A, Hayward JA, Lee J, Victor Makota F, Rug M, et al. A key cytosolic iron-sulfur cluster synthesis protein localises to the mitochondrion of Toxoplasma gondii. Mol Microbiol. 2020; mmi.14651. doi:10.1111/mmi.14651

21. Renaud EA, Maupin AJM, Besteiro S. Iron-sulfur cluster biogenesis and function in Apicomplexa parasites. Biochimica et Biophysica Acta (BBA) - Molecular Cell Research. 2025;1872: 119876. doi:10.1016/j.bbamcr.2024.119876

22. Renaud EA, Pamukcu S, Cerutti A, Berry L, Lemaire-Vieille C, Yamaryo-Botté Y, et al. Disrupting the plastidic iron-sulfur cluster biogenesis pathway in Toxoplasma gondii has pleiotropic effects irreversibly impacting parasite viability. J Biol Chem. 2022;298: 102243. doi:10.1016/j.jbc.2022.102243

23. Tsaousis AD, Gentekaki E, Eme L, Gaston D, Roger AJ. Evolution of the cytosolic iron-sulfur cluster assembly machinery in Blastocystis species and other microbial eukaryotes. Eukaryot Cell. 2014;13: 143–153. doi:10.1128/EC.00158-13

24. Pyrih J, Žárský V, Fellows JD, Grosche C, Wloga D, Striepen B, et al. The iron-sulfur scaffold protein HCF101 unveils the complexity of organellar evolution in SAR, Haptista and Cryptista. BMC Ecol Evol. 2021;21: 46. doi:10.1186/s12862-021-01777-x

25. Lezhneva L, Amann K, Meurer J. The universally conserved HCF101 protein is involved in assembly of [4Fe-4S]-cluster-containing complexes in *Arabidopsis thaliana* chloroplasts. The Plant Journal. 2004;37: 174–185. doi:10.1046/j.1365-313X.2003.01952.x

26. Schwenkert S, Netz DJA, Frazzon J, Pierik AJ, Bill E, Gross J, et al. Chloroplast HCF101 is a scaffold protein for [4Fe-4S] cluster assembly. Biochem J. 2009;425: 207–214. doi:10.1042/BJ20091290

27. Leipe DD, Wolf YI, Koonin EV, Aravind L. Classification and evolution of P-loop GTPases and related ATPases. Journal of Molecular Biology. 2002;317: 41–72. doi:10.1006/jmbi.2001.5378

28. Boyd JM, Drevland RM, Downs DM, Graham DE. Archaeal ApbC/Nbp35 homologs function as iron-sulfur cluster carrier proteins. J Bacteriol. 2009;191: 1490–1497. doi:10.1128/JB.01469-08

29. Mashruwala AA, Boyd JM. Investigating the role(s) of SufT and the domain of unknown function 59 (DUF59) in the maturation of iron–sulfur proteins. Curr Genet. 2018;64: 9–16. doi:10.1007/s00294-017-0716-5

30. Seeber F, Soldati-Favre D. Metabolic pathways in the apicoplast of apicomplexa. International Review of Cell and Molecular Biology. 2010. pp. 161–228.

31. Shrivastava D, Abboud E, Ramchandra JP, Jha A, Marq J-B, Chaurasia A, et al. ATM1, an essential conserved transporter in Apicomplexa, bridges mitochondrial and cytosolic [Fe-S] biogenesis. PLoS Pathog. 2024;20: e1012593. doi:10.1371/journal.ppat.1012593

32. Meurer J, Meierhoff K, Westhoff P. Isolation of high-chlorophyll-fluorescence mutants ofArabidopsis thaliana and their characterisation by spectroscopy, immunoblotting and Northern hybridisation. Planta. 1996;198: 385–396. doi:10.1007/BF00620055

33. Mathur V, Salomaki ED, Wakeman KC, Na I, Kwong WK, Kolisko M, et al. Reconstruction of plastid proteomes of apicomplexans and close relatives reveals the major evolutionary outcomes of cryptic plastids. Mol Biol Evol. 2023;40: msad002. doi:10.1093/molbev/msad002

34. Zhu G, Marchewka MJ, Keithly JS. Cryptosporidium parvum appears to lack a plastid genome. Microbiology. 2000;146: 315–321. doi:10.1099/00221287-146-2-315

35. Moore RB, Oborník M, Janouskovec J, Chrudimský T, Vancová M, Green DH, et al. A photosynthetic alveolate closely related to apicomplexan parasites. Nature. 2008;451: 959– 963. doi:10.1038/nature06635

36. Meissner M, Brecht S, Bujard H, Soldati D. Modulation of myosin A expression by a newly established tetracycline repressor-based inducible system in Toxoplasma gondii. Nucleic Acids Res. 2001;29: E115.

37. Cerutti A, Blanchard N, Besteiro S. The bradyzoite: a key developmental stage for the persistence and pathogenesis of toxoplasmosis. Pathogens. 2020;9: 234. doi:10.3390/pathogens9030234

38. Gubbels M-J, Wieffer M, Striepen B. Fluorescent protein tagging in Toxoplasma gondii: identification of a novel inner membrane complex component conserved among Apicomplexa. Molecular and Biochemical Parasitology. 2004;137: 99–110. doi:10.1016/j.molbiopara.2004.05.007

39. Francia ME, Striepen B. Cell division in apicomplexan parasites. Nat Rev Microbiol. 2014;12: 125–136. doi:10.1038/nrmicro3184

40. Maclean AE, Sloan MA, Renaud EA, Argyle BE, Lewis WH, Ovciarikova J, et al. The Toxoplasma gondii mitochondrial transporter ABCB7L is essential for the biogenesis of cytosolic and nuclear iron-sulfur cluster proteins and cytosolic translation. Weiss LM, editor. mBio. 2024; e00872–24. doi:10.1128/mbio.00872-24

41. Marquez MD, Greth C, Buzuk A, Liu Y, Blinn CM, Beller S, et al. Cytosolic iron–sulfur protein assembly system identifies clients by a C-terminal tripeptide. Proc Natl Acad Sci USA. 2023;120: e2311057120. doi:10.1073/pnas.2311057120

42. Nolan SJ, Romano JD, Kline JT, Coppens I. Novel approaches to kill Toxoplasma gondii by exploiting the uncontrolled uptake of unsaturated fatty acids and vulnerability to lipid storage inhibition of the parasite. Antimicrob Agents Chemother. 2018;62: e00347–18. doi:10.1128/AAC.00347-18

43. Jarc E, Petan T. Lipid droplets and the management of cellular stress. Yale J Biol Med. 2019;92: 435–452.

44. Nadipuram SM, Thind AC, Rayatpisheh S, Wohlschlegel JA, Bradley PJ. Proximity biotinylation reveals novel secreted dense granule proteins of Toxoplasma gondii bradyzoites. PLoS One. 2020;15: e0232552. doi:10.1371/journal.pone.0232552

45. Jung C, Lee CY-F, Grigg ME. The SRS superfamily of Toxoplasma surface proteins. International Journal for Parasitology. 2004;34: 285–296. doi:10.1016/j.ijpara.2003.12.004

46. Tomita T, Bzik DJ, Ma YF, Fox BA, Markillie LM, Taylor RC, et al. The Toxoplasma gondii cyst wall protein CST1 is critical for cyst wall integrity and promotes bradyzoite persistence. PLoS Pathog. 2013;9: e1003823. doi:10.1371/journal.ppat.1003823

47. Ben Chaabene R, Lentini G, Soldati-Favre D. Biogenesis and discharge of the rhoptries: Key organelles for entry and hijack of host cells by the Apicomplexa. Mol Microbiol. 2021;115: 453–465. doi:10.1111/mmi.14674

48. Mueller C, Klages N, Jacot D, Santos JM, Cabrera A, Gilberger TW, et al. The Toxoplasma protein ARO mediates the apical positioning of rhoptry organelles, a prerequisite for host cell invasion. Cell Host & Microbe. 2013;13: 289–301. doi:10.1016/j.chom.2013.02.001

49. Valasatava Y, Rosato A, Banci L, Andreini C. MetalPredator: a web server to predict iron-sulfur cluster binding proteomes. Bioinformatics. 2016;32: 2850–2852. doi:10.1093/bioinformatics/btw238

50. Barylyuk K, Koreny L, Ke H, Butterworth S, Crook OM, Lassadi I, et al. A comprehensive subcellular atlas of the Toxoplasma proteome via hyperLOPIT provides spatial context for protein functions. Cell Host & Microbe. 2020; S193131282030514X. doi:10.1016/j.chom.2020.09.011

51. Stehling O, Mascarenhas J, Vashisht AA, Sheftel AD, Niggemeyer B, Rösser R, et al. Human CIA2A-FAM96A and CIA2B-FAM96B integrate iron homeostasis and maturation of different Subsets of cytosolic-nuclear iron-sulfur proteins. Cell Metabolism. 2013;18: 187–198. doi:10.1016/j.cmet.2013.06.015

52. Schmidt EK, Clavarino G, Ceppi M, Pierre P. SUnSET, a nonradioactive method to monitor protein synthesis. Nat Methods. 2009;6: 275–277. doi:10.1038/nmeth.1314

53. Bych K, Kerscher S, Netz DJA, Pierik AJ, Zwicker K, Huynen MA, et al. The iron–sulphur protein Ind1 is required for effective complex I assembly. EMBO J. 2008;27: 1736–1746. doi:10.1038/emboj.2008.98

54. Aubert C, Mandin P, Py B. Mrp and SufT, two bacterial homologs of eukaryotic CIA Factors involved in Fe-S clusters biogenesis. Inorganics. 2023;11: 431. doi:10.3390/inorganics11110431

55. Stehling O, Jeoung J-H, Freibert SA, Paul VD, Bänfer S, Niggemeyer B, et al. Function and crystal structure of the dimeric P-loop ATPase CFD1 coordinating an exposed [4Fe-4S] cluster for transfer to apoproteins. Proc Natl Acad Sci U S A. 2018;115: E9085–E9094. doi:10.1073/pnas.1807762115

56. Mashruwala AA, Bhatt S, Poudel S, Boyd ES, Boyd JM. The DUF59 containing protein SufT is involved in the maturation of Iron-Sulfur (FeS) proteins during conditions of high FeS cofactor demand in Staphylococcus aureus. PLoS Genet. 2016;12: e1006233. doi:10.1371/journal.pgen.1006233

57. Sidik SM, Huet D, Ganesan SM, Huynh M-H, Wang T, Nasamu AS, et al. A Genome-wide CRISPR Screen in Toxoplasma Identifies Essential Apicomplexan Genes. Cell. 2016;166: 1423–1435.e12. doi:10.1016/j.cell.2016.08.019

58. Liang D, Minikes AM, Jiang X. Ferroptosis at the intersection of lipid metabolism and cellular signaling. Molecular Cell. 2022;82: 2215–2227. doi:10.1016/j.molcel.2022.03.022

59. Renaud EA, Maupin AJM, Bordat Y, Graindorge A, Berry L, Besteiro S. Iron depletion has different consequences on the growth and survival of Toxoplasma gondii strains. Virulence. 2024;15: 2329566. doi:10.1080/21505594.2024.2329566

60. Paul VD, Mühlenhoff U, Stümpfig M, Seebacher J, Kugler KG, Renicke C, et al. The deca-GX3 proteins Yae1-Lto1 function as adaptors recruiting the ABC protein Rli1 for iron-sulfur cluster insertion. eLife. 2015;4: e08231. doi:10.7554/eLife.08231

61. Zhai C, Li Y, Mascarenhas C, Lin Q, Li K, Vyrides I, et al. The function of ORAOV1/LTO1, a gene that is overexpressed frequently in cancer: essential roles in the function and biogenesis of the ribosome. Oncogene. 2014;33: 484–494. doi:10.1038/onc.2012.604

62. Prusty NR, Camponeschi F, Ciofi-Baffoni S, Banci L. The human YAE1-ORAOV1 complex of the cytosolic iron-sulfur protein assembly machinery binds a [4Fe-4S] cluster. Inorganica Chimica Acta. 2021;518: 120252. doi:10.1016/j.ica.2021.120252

63. Navarro-Quiles C, Mateo-Bonmatí E, Micol JL. ABCE Proteins: From Molecules to development. Front Plant Sci. 2018;9: 1125. doi:10.3389/fpls.2018.01125

64. Sheiner L, Demerly JL, Poulsen N, Beatty WL, Lucas O, Behnke MS, et al. A Systematic Screen to Discover and Analyze Apicoplast Proteins Identifies a Conserved and Essential Protein Import Factor. PLoS Pathog. 2011;7. doi:10.1371/journal.ppat.1002392

65. Grosche C, Diehl A, Rensing SA, Maier UG. Iron–sulfur cluster biosynthesis in algae with complex plastids. Embley M, editor. Genome Biology and Evolution. 2018;10: 2061–2071. doi:10.1093/gbe/evy156

66. Katoh K, Rozewicki J, Yamada KD. MAFFT online service: multiple sequence alignment, interactive sequence choice and visualization. Briefings in Bioinformatics. 2019;20: 1160–1166. doi:10.1093/bib/bbx108

67. Criscuolo A, Gribaldo S. BMGE (Block Mapping and Gathering with Entropy): a new software for selection of phylogenetic informative regions from multiple sequence alignments. BMC Evol Biol. 2010;10: 210. doi:10.1186/1471-2148-10-210

68. Trifinopoulos J, Nguyen L-T, von Haeseler A, Minh BQ. W-IQ-TREE: a fast online phylogenetic tool for maximum likelihood analysis. Nucleic Acids Res. 2016;44: W232–W235. doi:10.1093/nar/gkw256

69. Minh BQ, Nguyen MAT, Von Haeseler A. Ultrafast approximation for phylogenetic bootstrap. Molecular Biology and Evolution. 2013;30: 1188–1195. doi:10.1093/molbev/mst024

70. Tamura K, Stecher G, Kumar S. MEGA11: Molecular Evolutionary Genetics Analysis version 11. Battistuzzi FU, editor. Molecular Biology and Evolution. 2021;38: 3022–3027. doi:10.1093/molbev/msab120

71. Letunic I, Bork P. Interactive Tree of Life (iTOL) v6: recent updates to the phylogenetic tree display and annotation tool. Nucleic Acids Research. 2024;52: W78–W82. doi:10.1093/nar/gkae268

72. Lévêque MF, Berry L, Cipriano MJ, Nguyen H-M, Striepen B, Besteiro S. Autophagy-related protein ATG8 has a noncanonical function for apicoplast inheritance in Toxoplasma gondii. MBio. 2015;6: e01446–15.

73. Nguyen HM, Liu S, Daher W, Tan F, Besteiro S. Characterisation of two Toxoplasma PROPPINs homologous to Atg18/WIPI suggests they have evolved distinct specialised functions. PLoS ONE. 2018;13: e0195921. doi:10.1371/journal.pone.0195921

74. Kong-Hap MA, Mouammine A, Daher W, Berry L, Lebrun M, Dubremetz J-F, et al. Regulation of ATG8 membrane association by ATG4 in the parasitic protist Toxoplasma gondii. Autophagy. 2013;9: 1334–1348. doi:10.4161/auto.25189

75. Anderson-White BR, Ivey FD, Cheng K, Szatanek T, Lorestani A, Beckers CJ, et al. A family of intermediate filament-like proteins is sequentially assembled into the cytoskeleton of Toxoplasma gondii. Cell Microbiol. 2011;13: 18–31. doi:10.1111/j.1462-5822.2010.01514.x

76. Couvreur G, Sadak A, Fortier B, Dubremetz JF. Surface antigens of Toxoplasma gondii. Parasitology. 1988;97 (Pt 1): 1–10.

77. Herm-Gotz A. Toxoplasma gondii myosin A and its light chain: a fast, single-headed, plus-end-directed motor. The EMBO Journal. 2002;21: 2149–2158. doi:10.1093/emboj/21.9.2149

78. Hajj HE, Lebrun M, Fourmaux MN, Vial H, Dubremetz JF. Characterization, biosynthesis and fate of ROP7, a ROP2 related rhoptry protein of Toxoplasma gondii⋆. Molecular and Biochemical Parasitology. 2006;146: 98–100. doi:10.1016/j.molbiopara.2005.10.011

79. Crawford MJ, Thomsen-Zieger N, Ray M, Schachtner J, Roos DS, Seeber F. Toxoplasma gondii scavenges host-derived lipoic acid despite its de novo synthesis in the apicoplast. EMBO J. 2006;25: 3214–3222. doi:10.1038/sj.emboj.7601189

80. Berger N, Vignols F, Przybyla-Toscano J, Roland M, Rofidal V, Touraine B, et al. Identification of client iron–sulfur proteins of the chloroplastic NFU2 transfer protein in Arabidopsis thaliana. Takahashi H, editor. Journal of Experimental Botany. 2020;71: 4171–4187. doi:10.1093/jxb/eraa166

81. Cox J, Mann M. MaxQuant enables high peptide identification rates, individualized p.p.b.-range mass accuracies and proteome-wide protein quantification. Nat Biotechnol. 2008;26: 1367– 1372. doi:10.1038/nbt.1511

82. Hehl AB, Basso WU, Lippuner C, Ramakrishnan C, Okoniewski M, Walker RA, et al. Asexual expansion of Toxoplasma gondii merozoites is distinct from tachyzoites and entails expression of non-overlapping gene families to attach, invade, and replicate within feline enterocytes. BMC Genomics. 2015;16: 66. doi:10.1186/s12864-015-1225-x

83. Smyth GK. Linear models and empirical bayes methods for assessing differential expression in microarray experiments. Stat Appl Genet Mol Biol. 2004;3: Article3. doi:10.2202/1544-6115.1027

84. Daniel Gietz R, Woods RA. Transformation of yeast by lithium acetate/single-stranded carrier DNA/polyethylene glycol method. Methods in Enzymology. Elsevier; 2002. pp. 87–96. doi:10.1016/S0076-6879(02)50957-5

85. Semenovskaya K, Lévêque MF, Berry L, Bordat Y, Dubremetz J, Lebrun M, et al. TgZFP2 is a novel zinc finger protein involved in coordinating mitosis and budding in Toxoplasma. Cellular Microbiology. 2020;22: e13120.

86. Henkel S, Frohnecke N, Maus D, McConville MJ, Laue M, Blume M, et al. Toxoplasma gondii apicoplast-resident ferredoxin is an essential electron transfer protein for the MEP isoprenoid-biosynthetic pathway. J Biol Chem. 2022;298: 101468. doi:10.1016/j.jbc.2021.101468

87. Lamarque MH, Roques M, Kong-Hap M, Tonkin ML, Rugarabamu G, Marq J-B, et al. Plasticity and redundancy among AMA-RON pairs ensure host cell entry of Toxoplasma parasites. Nat Commun. 2014;5: 4098. doi:10.1038/ncomms5098

