## Supplemental Figures S1 to S12 for "HCF101 is a novel component of the CIA cytosolic iron-sulfur synthesis pathway in the human pathogen *Toxoplasma gondii*"

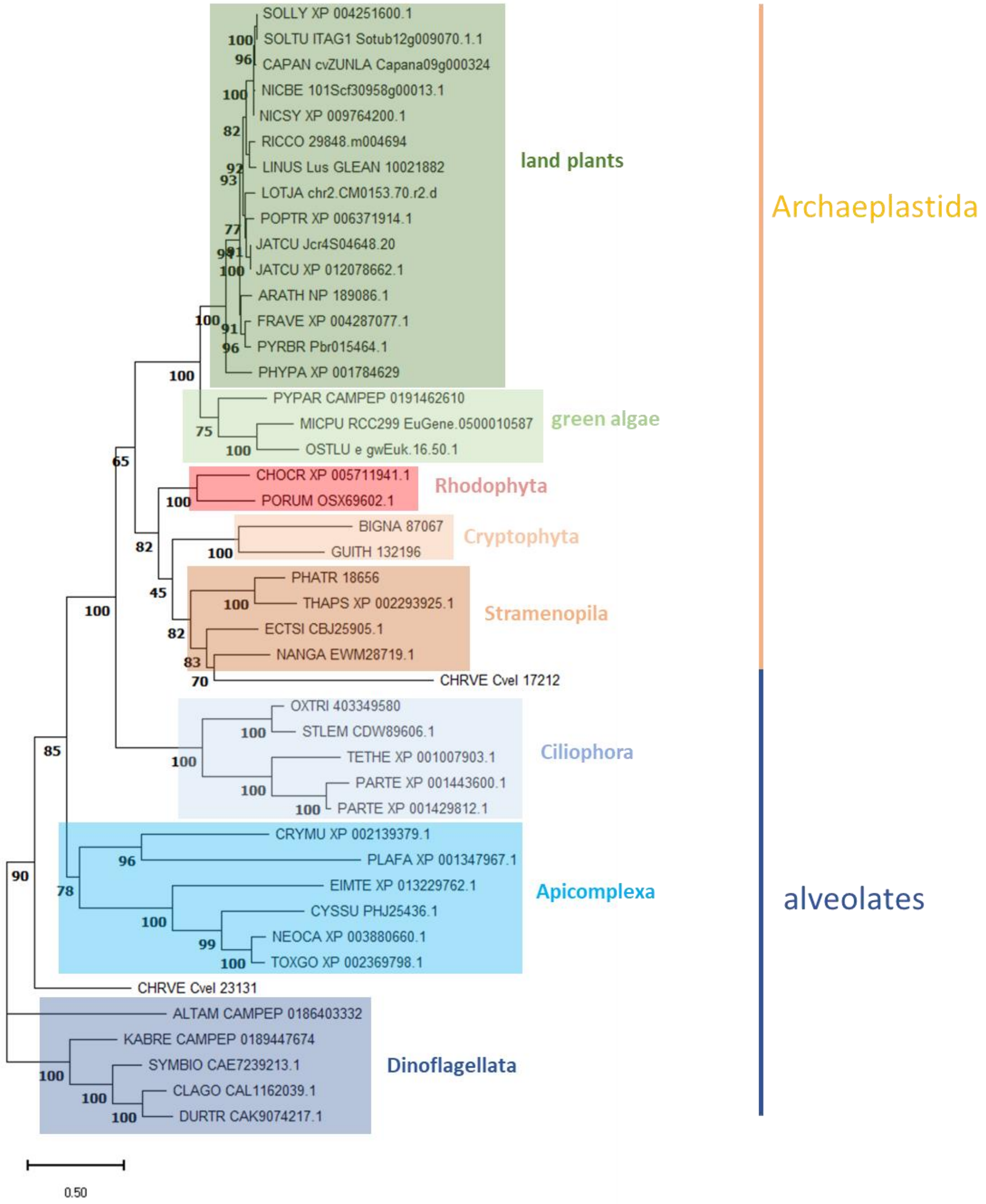


**S1 Figure. Phylogenetic analysis of selected HCF101 homologs.** Unrooted tree was generated by aligning sequences from HCF101 homologs the resulting alignment was submitted to phylogenetic analysis with the maximum likelihood method. Bootstrap values, inferred from 500 replicates, are indicated at the nodes. Scale bar represents 0.5 residue substitution per site.

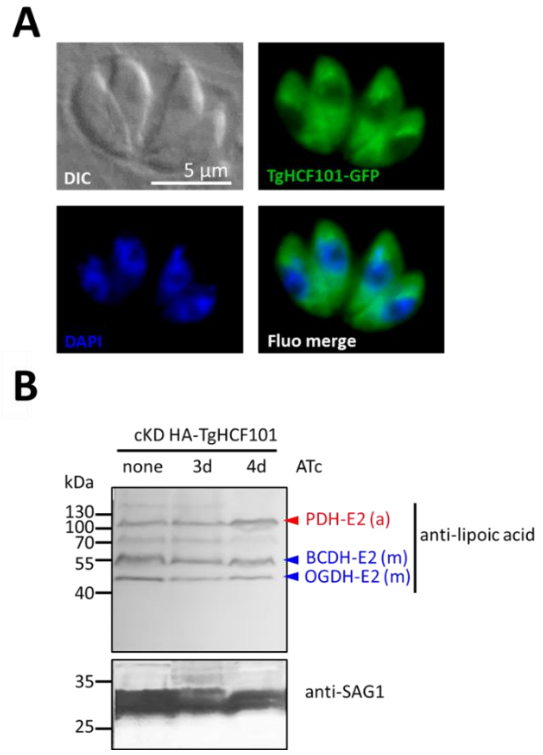


**S2 Figure. TgHCF101 is not associated with the apicoplast or its metabolism. A.** Fluorescent imaging of *T. gondii* tachyzoites ectopically-expressing a GFP-fused copy of TgHCF101, DNA was stained with DAPI. Scale bar= 5 µm. **B.** Immunoblot analysis of the lipoylation profile of *T. gondii* tachyzoites, typically showing apicoplast (Pyruvate dehydrogenase subunit E2, PDH-E2) and mitochondrial (Branched-chain 2-oxo acid dehydrogenase, BCDH-E2 and 2-oxoglutarate dehydrogenase, OGDH-E2) proteins that are largely unaffected upon TgHCF101 depletion by incubation with ATc for up to 4 days. Anti-SAG1 antibody is used as a loading control.

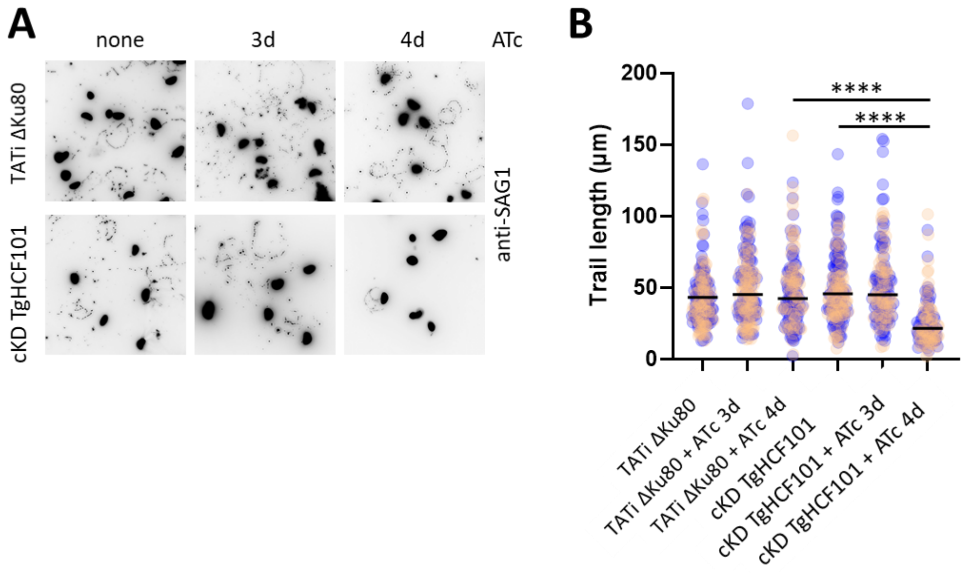


**S3 Figure. TgHCF101 depletion does not initially impact parasite motility. A.** Representative images of a gliding motility assay selected out of 3 independent replicates. The trails left behind by gliding parasites were detected using an anti-SAG1 antibody. TgHCF101 conditional mutant and parental cell line (TATi ΔKu80) were grown for 3 or 4 days in the presence of ATc. **B.** Quantification of SAG1 trail length showed in (A) produced by parasites measured on 10 randomly selected fields. At least 100 trails were measured for each dataset, values represented are means of n=2 independent biological replicates (red and blue circles represent datapoints from these replicates), **** p-value≤0.0001, Student's *t*-test.

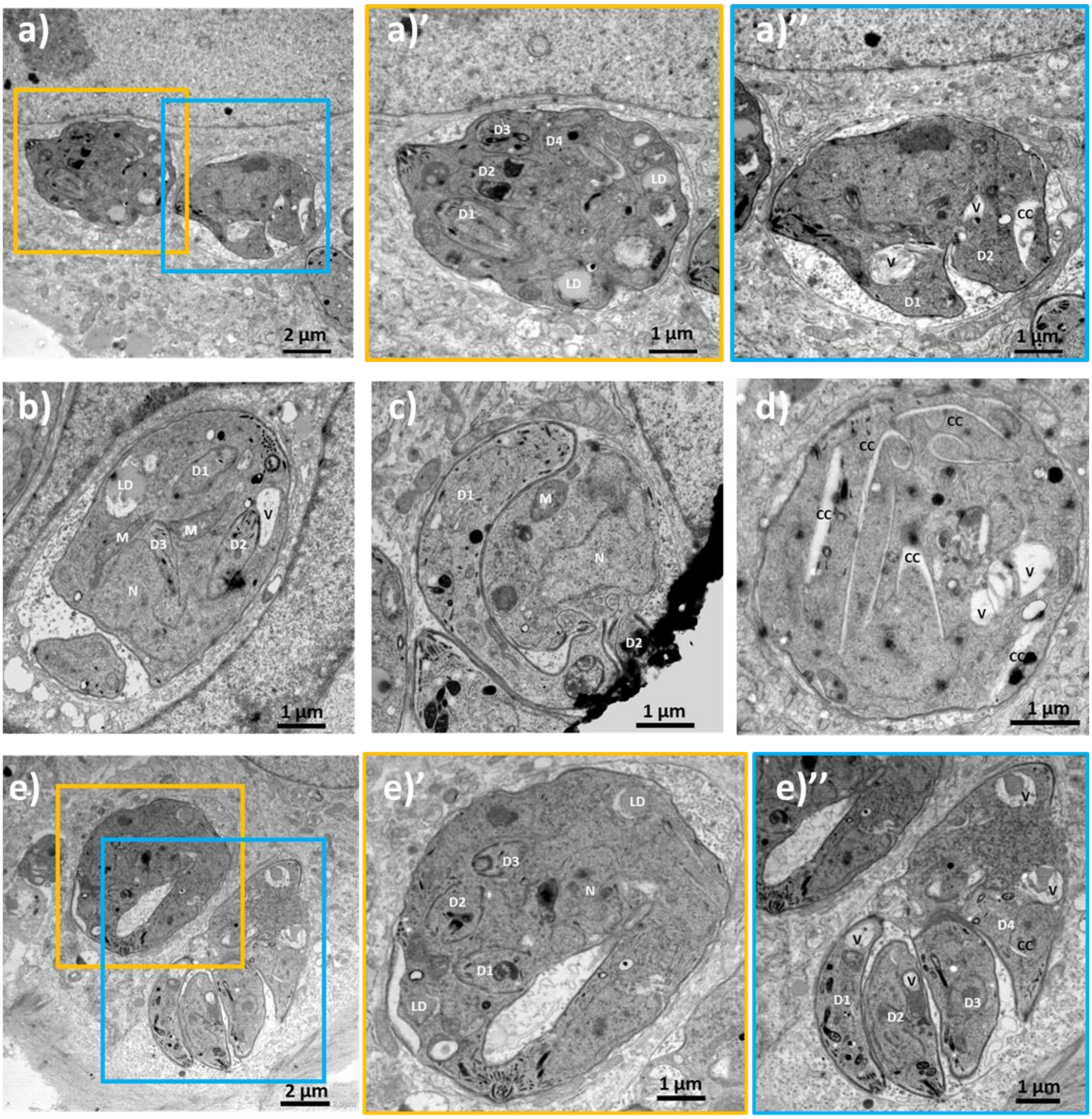


**S4 Figure. Additional examples of morphological defects caused by TgHCF101 depletion.** Electron microscopy was performed on cKD-TgHCF101 parasites pre-incubated with ATc for 24h before being released from their host cell and allowed to reinvade for 24h in the presence (+ATc). CC: cytoplasmic cleft, D: daughter bud, LD: lipid droplet, M: mitochondrion, N: nucleus, V: vacuole. ' and " denote magnifications of different regions of the same lettered image.

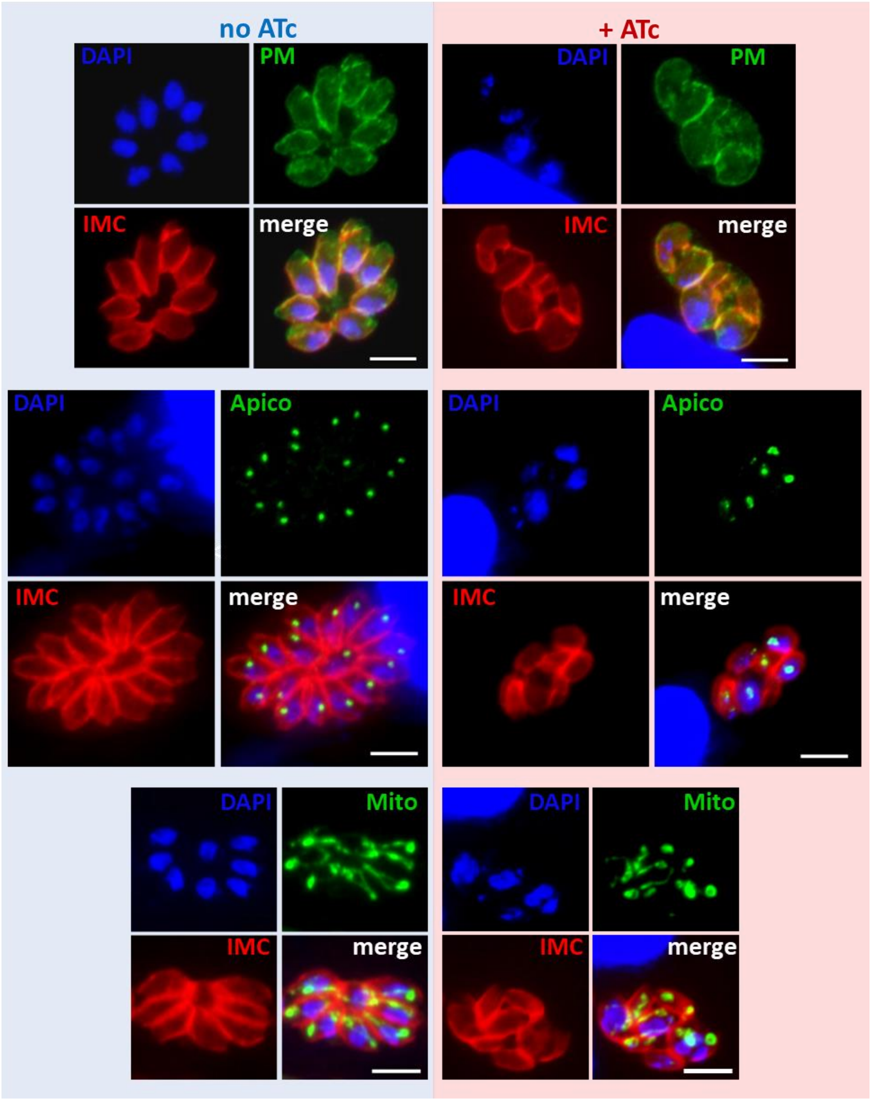


**S5 Figure. Immunofluorescence analysis of cKD HA-TgHCF101 parasites incubated or not with ATc for 4 days, showing organellar defects upon TgHCF101 depletion.** Mito: mitochondrion, labeled with an anti-F1β ATPAse antibody; Apico: apicoplast, labeled with an anti-PDH-E2 antibody; IMC: inner membrane complex, labeled with an anti-IMC3 antibody; PM: plasma membrane, labeled with an anti-SAG1 antibody; DNA was labeled with DAPI. Scale bar = 5 µm.

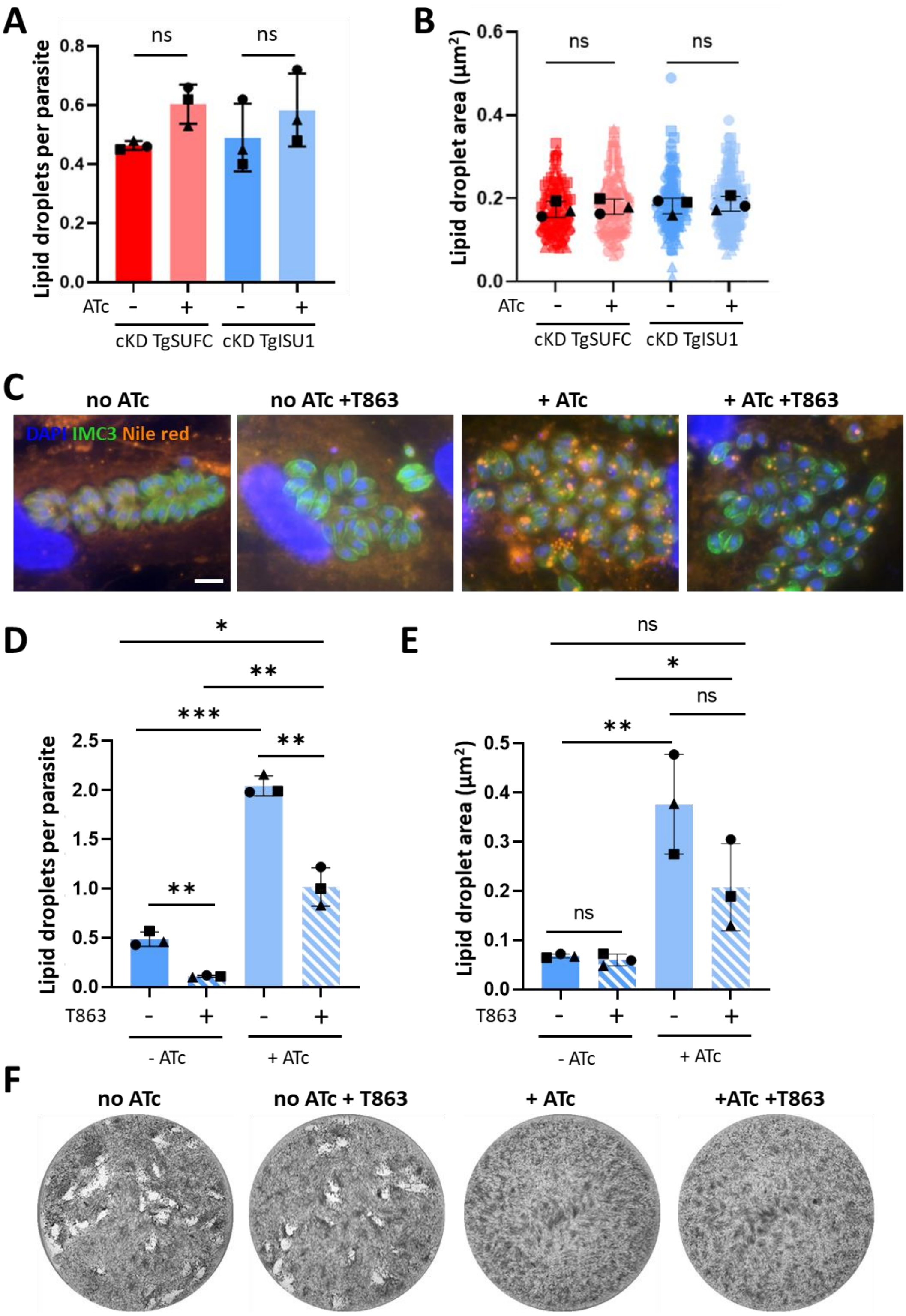


**S6 Fig. Lipid droplet induction depends on TgHCF101 depletion and is not responsible for parasite demise. A.** and **B.** correspond to quantification of the number and area of lipid droplets, respectively, in apicoplast (cKD-TgSUFC) and mitochondrion (cKD-TgISU1) Fe-S cluster synthesis mutants grown in the absence or presence of ATc for up to 72h. 100 parasites were analyzed per condition. Values are the mean ± SD of n=3 independent biological replicates; ns, not significant (p-value ≥ 0.05, Student's *t*-test). **C.** Immunofluorescence assay of parasites from the cKD-TgHCF101 cell line treated for 72h with or in the absence of ATc and supplemented or not with diacylglycerol acyltransferase inhibitor T863.

Lipid droplets were detected with Nile red (orange), parasites are outlined with anti-IMC3 antibody (green) and DNA is stained with DAPI. Scale bar= 5 µm. For this experiment and the following analyses involving T863, the drug was used at 5 µM and an equivalent volume of the vehicle only (DMSO) was used in the control conditions. **D.** and **E.** correspond to the quantification of the number and area of lipid droplet, 100 parasites were analyzed per condition. Parasites were grown in the absence of ATc for the parental (TATi ΔKu80) and transgenic (cKD HA-TgHCF101) cell lines or for 72h with ATc treatment, supplemented or not with T863. Values are represented as the mean ± SD of n=3 independent biological replicates; ns, not significant (p-value ≥ 0.05), * p-value ≤ 0.05, ** p-value ≤ 0.01 and *** p-value < 0.001, Student's *t*-test. **F.** Plaque assays were carried out by infecting a monolayer of HFFs with cKD HA-TgHCF101 cell lines for 7 days in the presence or absence of ATc and supplemented or not with T863.

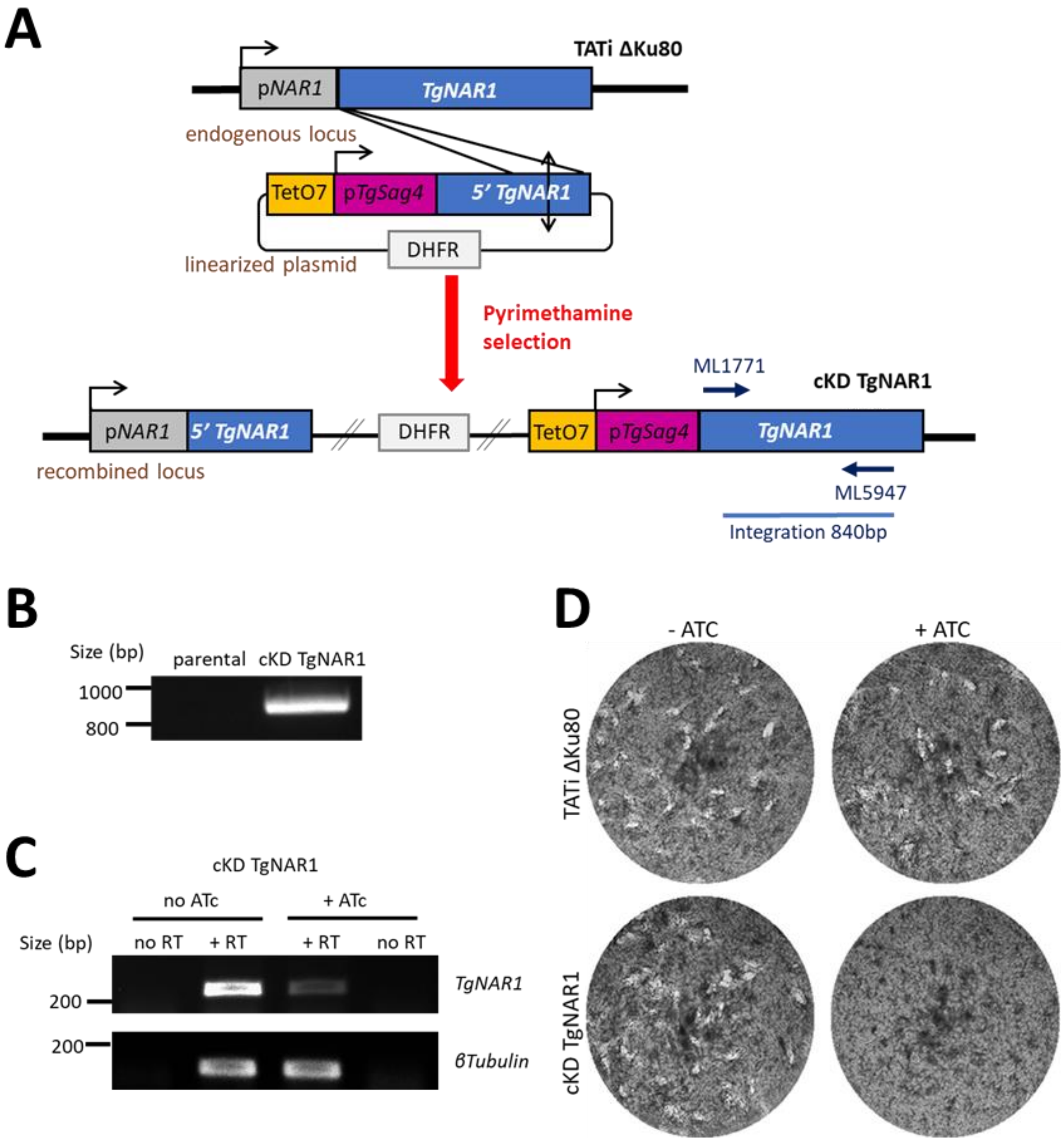


**S7 Figure. Generation of a TgNAR1 conditional mutant. A.** Strategy for generating the inducible knockdown of *TgNAR1* by promoter replacement in the TATi ΔKu80 cell line. **B.** Diagnostic PCR for checking correct integration of using the primers mentioned in **A**), on genomic DNAs of a transgenic parasite clone and of the parental strain. **C.** Semi-quantitative RT-PCR was used to verify the efficient down-regulation of *TgNAR1* upon addition of ATc. Reverse transcriptase was omitted in the 'no RT' control. *β-tubulin* was used as a control. **D.** Plaque assay revealed that depletion of TgNAR1 is important for parasite growth.

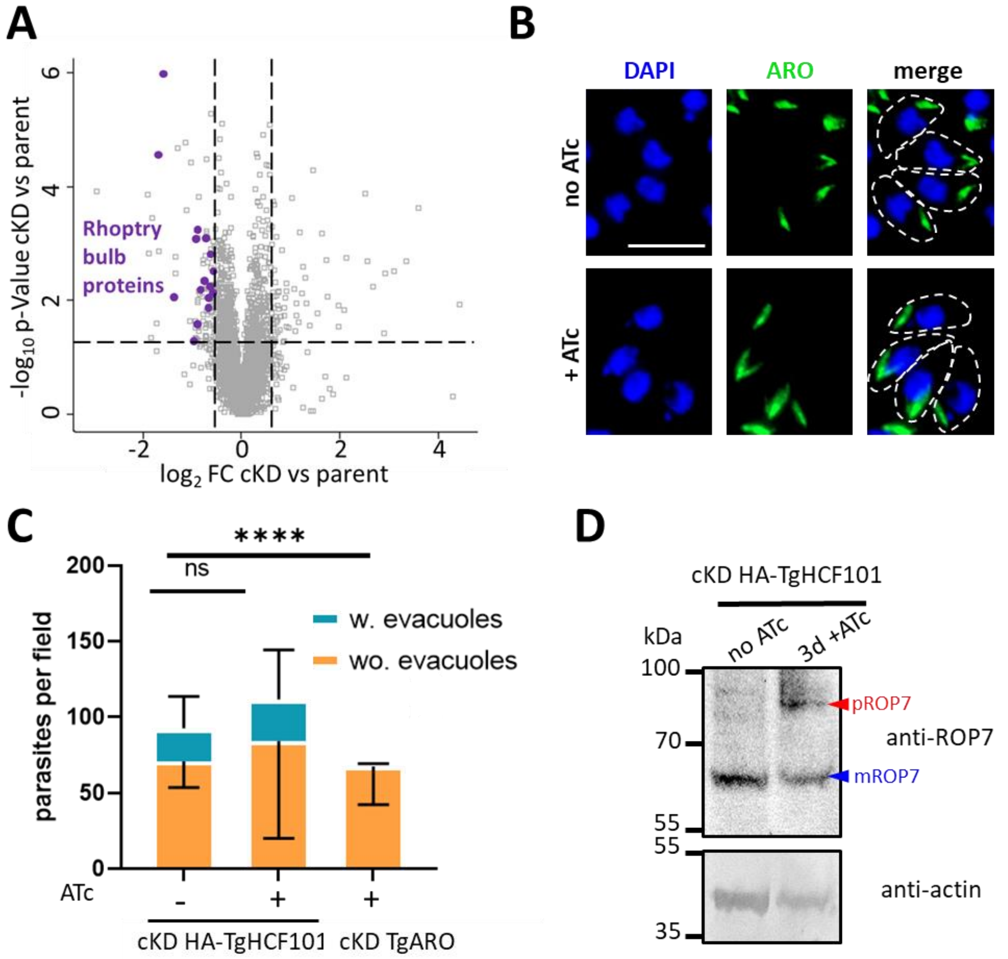


**S8 Figure. Depletion of TgHCF101 has no extensive impact on rhoptries. A.** Volcano plot showing differential expression of proteins impacted by TgHCF101 depletion after 72h of ATc treatment analyzed by label free quantitative proteomic. X-axis correspond to the $\log_2$ of the Fold-change, Y-axis correspond to the $-\log_{10}$ of the p-value when comparing cKD-TgHCF101 expression values to the TATi ΔKu80 parental cell line. Statistical analyses were performed with ANOVA from 4 independent biological replicates. Cut-offs were set at ≤1.5- or ≥1.5-FC and p-value ≤0.05. Significant hits corresponding to rhoptry bulb protein proteins were highlighted in purple on the graph. **B.** Immunofluorescence assay of parasites from the cKD HA-TgHCF101 cell line pre-treated for 48h and allowed to grow on HFF coated coverslips for another 24h with or in the absence of ATc. Rhoptries were detected with anti-ARO antibody (green), parasites are outlined with white dotted lines and DNA is stained with DAPI. Scale bar= 5µm**. C.** Quantification of rhoptry secretion events (evacuoles) in the cKD HA-TgHCF101 mutant upon TgHCF101 depletion for 72h. The TgARO conditional knock-down cell line serves as a control of rhoptry secretion defect upon ATc treatment. Parasites with and without evacuoles were counted on 20 randomly selected fields, with 3 technical replicates for each biological replicate. Values are represented as the mean ± SD of n=3 independent biological replicates; ns, not significant (p-value ≥ 0.05), * p-value ≤ 0.05, **** a p-value ≤ 0.001, Student's *t*-test. **D.** Immunoblot analysis of the expression of protein TgROP7 upon depletion of TgHCF101 for 3 days. The pro-form of TgROP7 (pROP7) is highlighted by a red arrow and the mature form (mROP7) by a blue arrow. TgROP7 signal was detected by anti-ROP7 antibody and anti-actin antibody was used as a loading control.

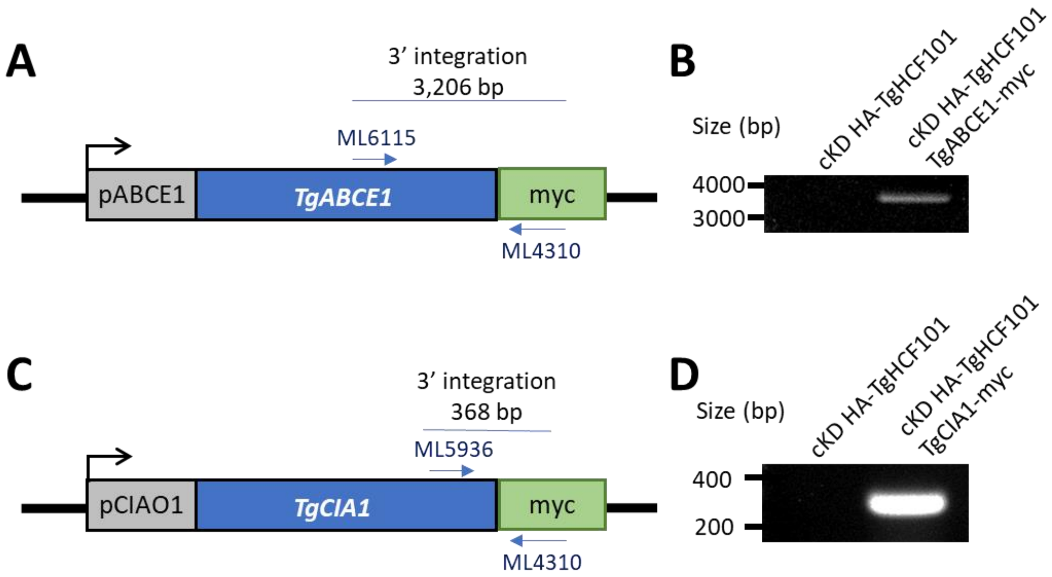


**S9 Figure. Constructs for tagging TgABCE1 and TgCIA1 in the cKD HA-TgHCF101 mutant background.** **A.** and **C.** correspond to schematic representations of the strategy used to add a C-terminal myc tag to proteins TgABCE1 (A) and TgCIA1 (C) by homologous recombination at the native locus. Chloramphenicol was used to select transgenic parasites. **B.** and **D.** correspond to diagnostic PCRs on genomic DNA from parental cell line (cKD HA-TgHCF101) or new clonal cell lines, in order to check for the integration of the sequence coding for the myc tag using primers highlighted on the (A) and (C) schemes.

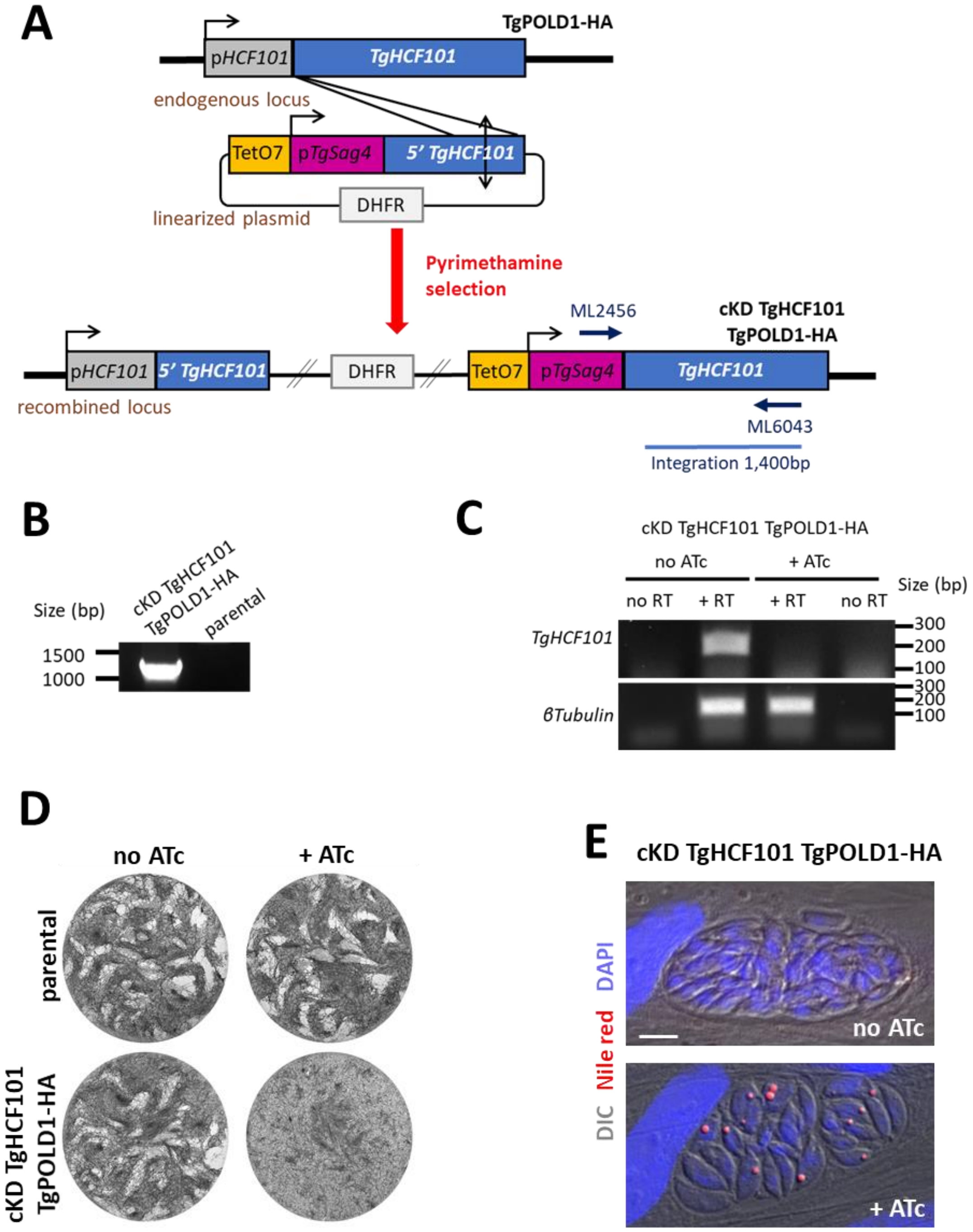


**S10 Figure. Generating a conditionally TgHCF101 mutant in the TgPOLD1-HA cell line. A.** Strategy for generating the inducible knockdown of *TgHCF101* by promoter replacement in the TgPOLD1-HA cell line. **B.** Diagnostic PCR for checking correct integration of using the primers mentioned in **A**), on genomic DNAs of a transgenic parasite clone and of the parental strain. **C.** Semi-quantitative RT-PCR was used to verify the efficient down-regulation of *TgHCF101* upon addition of ATc. Reverse transcriptase was omitted in the 'no RT' control. *β-tubulin* was used as a control. **D.** Plaque assay confirmed that depletion of TgHCF101 is important for parasite growth. **E.** Nile red staining confirmed that depletion of TgHCF101 in the cKD TgHCF101 TgPOLD1-HA cell line induces lipid droplet formation.

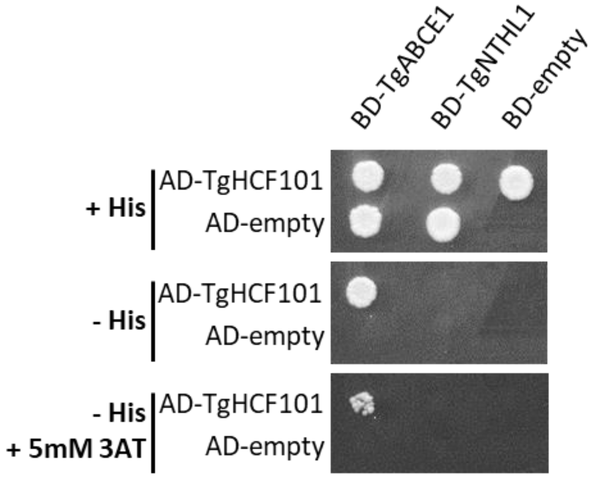


**S11 Figure. No direct interaction between TgHCF101 and TgNTHL1 was detected in a Gal4-based yeast two-hybrid assay.** Co-transformed YRG2 cells expressing AD- and BD-fusion proteins were plated on a control plate (+His, upper panel) for checking cell fitness and on the Y2H test plate (-His, mid panel), and plates were incubated at 30°C. Yeast growth was recorder after 5 days. At the cell concentration used (OD600 =0.05), the observed interaction between AD-TgHCF101 and BD-TgABCE1 fusion proteins was strong enough to allow cell growth in the presence of 5 mM 3AT inhibitor (-his + 3AT lower panel) but no interaction was detected with TgNTHL1. None of the proteins tested alone exhibited HIS3 transactivation capacities. Results shown here are representative of three independent experiments.

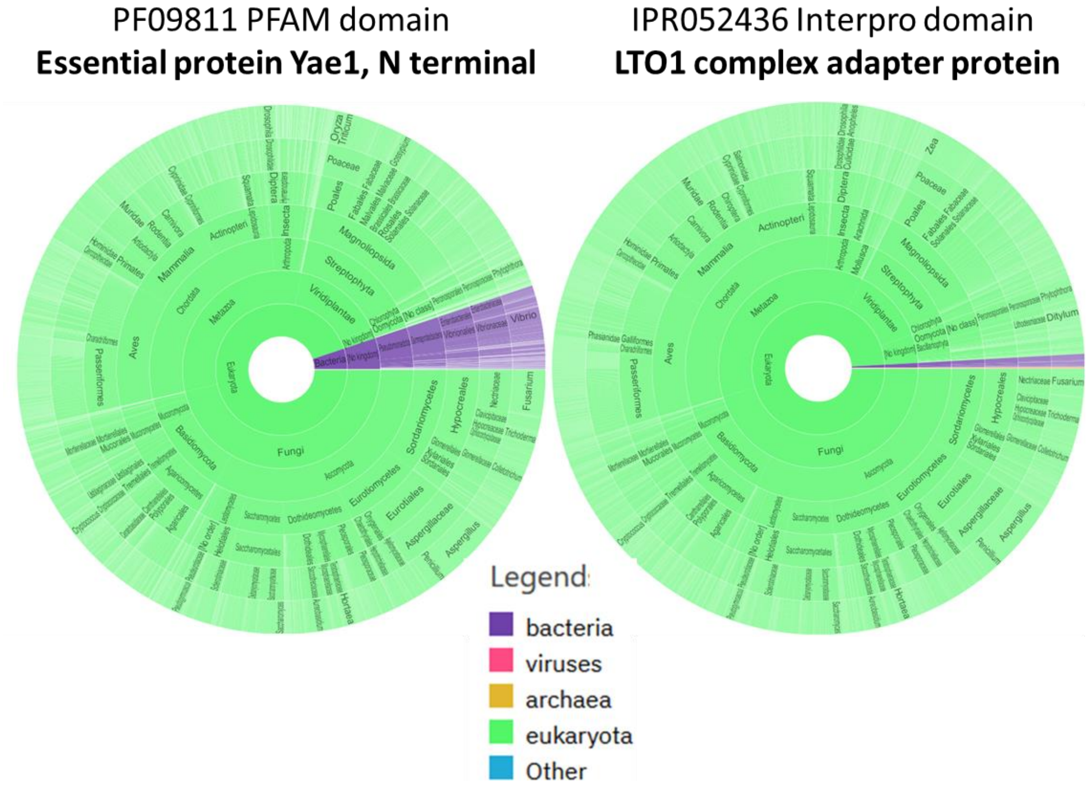


**S12 Figure. CIA adapters YAE1 and LTO1 are restricted to specific eukaryotic lineages.** Sunburst representation of the distribution of YAE1 and LTO1 domains as retrieved in the Interpro database (https://www.ebi.ac.uk/interpro/).
